# Reciprocal interactions between EMT and BMP signalling drive collective cell invasion

**DOI:** 10.64898/2025.12.02.691808

**Authors:** Yuri Takahashi, Alexandra Neaverson, Lara Busby, Filip Twarowski, Carlos Camacho-Macorra, Guillermo Serrano Nájera, Benjamin Steventon

## Abstract

During collective cell invasion, epithelial-to-mesenchymal transition (EMT) and morphogen signalling-mediated cell fate specification are traditionally viewed as a linear cascade: morphogens drive cell fates that activate EMT programs. Here, we uncover reciprocal coupling between EMT initiation and BMP signalling mediated by *SNAI2* and *SMAD1*. Using substrate-induced EMT in *ex vivo* explants, we demonstrate that EMT initiation upregulates *SMAD1* expression, priming cells for BMP signalling competence across germ layers. Single-cell RNA sequencing reveals *SNAI2* and *SMAD1* co-expression in EMT initiation regions, and *SNAI2* overexpression is sufficient to induce ectopic *SMAD1* expression *in vivo*. While BMP signalling is dispensable for EMT initiation, it regulates cell fate proportions, dispersal dynamics, precursor region depletion rates, and migration directionality. This coupling provides a mechanism for synchronising cell fate specification with invasion progression during axis elongation, positioning EMT as a process that actively modulates morphogen competence to coordinate tissue-level cell behaviours during collective cell invasion.

**Highlights:**

- EMT initiation directly primes BMP pathway competence.
- *SNAI2* overexpression is sufficient to drive ectopic *SMAD1* expression.
- BMP signalling is spatiotemporally restricted to EMT initiation zones.
- BMP signalling tunes cell fate proportions and invasion dynamics.

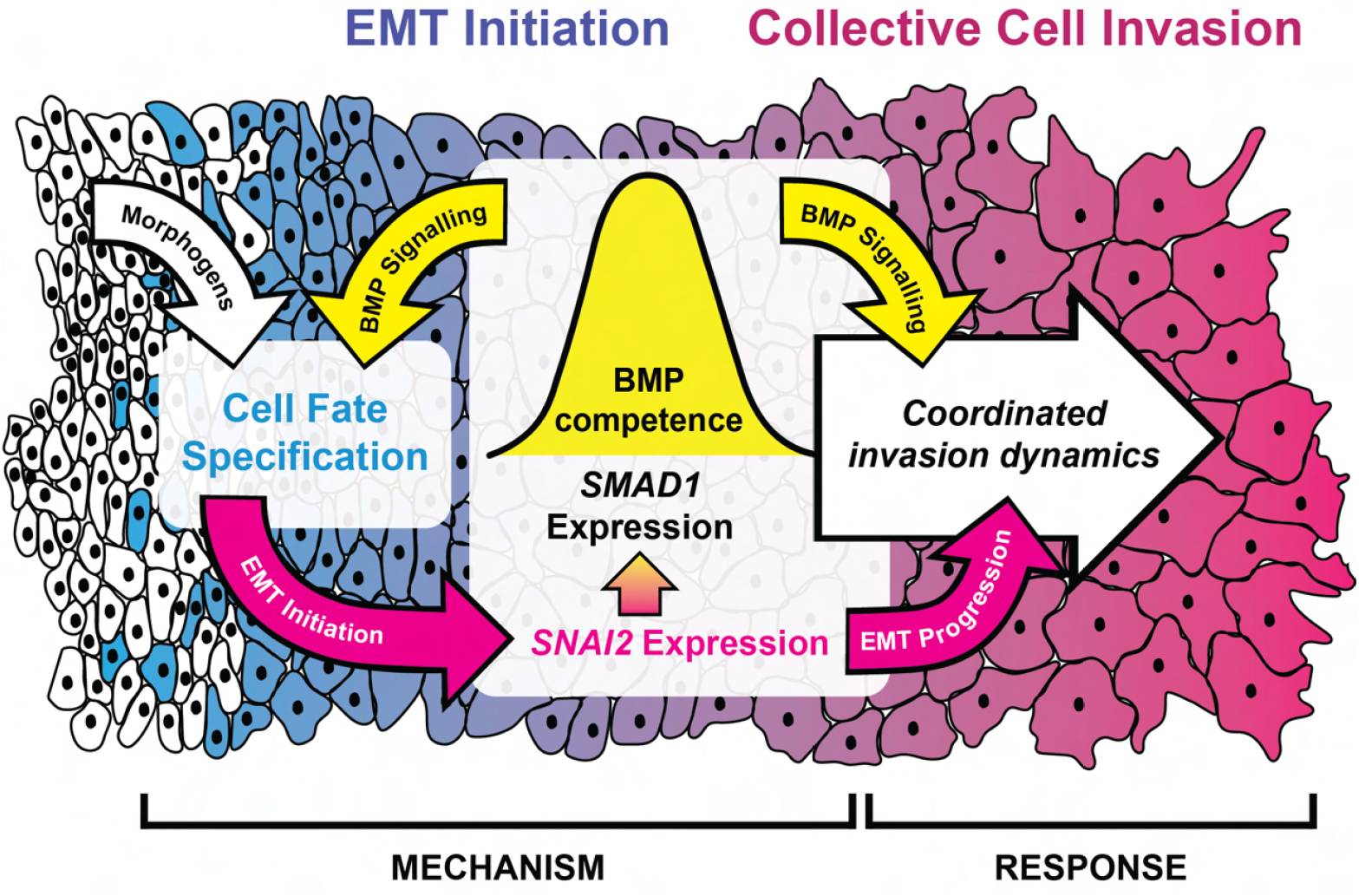

## Introduction

Collective cell invasion is the coordinated multi-step process where groups of cells penetrate tissue boundaries and move through complex environments. This process encompasses ingression, delamination, and migration, therefore requiring tight spatiotemporal coordination of cell behaviours. Two critical examples occur during early vertebrate development: mesoderm ingression through the primitive streak during gastrulation (reviewed in Chuai et al., 2012; Tam and Behringer, 1997; see also Voiculescu et al., 2014), which continues throughout axis elongation, and neural crest cell delamination during neurulation (reviewed in Theveneau and Mayor, 2012; see also Theveneau et al., 2010). In both contexts, populations of cells must simultaneously undergo epithelial-to-mesenchymal transition (EMT) while specifying appropriate migratory cell fates—a coordination challenge that demands precise integration of morphogen signalling with cellular state transitions.

Our current understanding of how collective cell invasion is regulated is based on a linear cascade model in which morphogens act upstream of cell fate specification and EMT (reviewed in Acloque et al., 2009; Thiery et al., 2009; Leathers and Rogers, 2022; Zhang et al., 2025). Morphogens such as Wnts, FGFs, and BMPs first specify cell fates that are primed to undergo migration as part of their differentiation programme. These fate-specified cells then activate transcriptional programs (driven by key EMT drivers such as the SNAIL and ZEB family transcription factors) that repress epithelial gene programs and activate mesenchymal ones that enable cells to lose epithelial polarity and remodel the surrounding extracellular matrix (reviewed in Debnath et al., 2022; see also Duong and Erickson, 2004; Brown et al., 2021). However, this linear cascade does not easily explain the tight population-level spatiotemporal coordination observed during collective cell invasion. Recent work has revealed that self-organized collective cell behaviours can emerge from reciprocal interactions between cellular state transitions and morphogen signalling, in which cellular behaviours both generate and respond to tissue-scale properties (Pfeifer et al., 2024). This raises a fundamental question: do reciprocal interactions exist between EMT initiation and morphogen signalling to drive collective cell invasion?

To investigate potential reciprocal coupling between EMT initiation and morphogen signalling, we employed an experimental system to induce EMT independently of developmental context. We adapted a chick embryo explant assay (Sanders, 1980; Newgreen, 1984; Busby et al., 2024) where mesodermal tissues are dissected and plated on extracellular matrix (ECM)-coated glass to assess migratory behaviour. We used this assay to compare two gastrula-stage chick embryonic tissues of different endogenous EMT states: the anterior primitive streak (APS) and the prospective neural plate (PNP). In both populations, *SNAI2* expression increased together with BMP pathway activation and subsequent BMP-induced cell fate changes in both germ layers. *In vivo* functional studies demonstrated that *SNAI2* overexpression is sufficient to induce ectopic *SMAD1* expression. We find that BMP signalling was dispensable for EMT initiation *in vivo*, prompting the question: does BMP signalling regulate the dynamics of collective cell invasion itself? To address this, we turned to *in vivo* grafting experiments that revealed that BMP signalling modulates cell dispersal dynamics, precursor region depletion rates, and migration directionality. Taken together, these findings reveal reciprocal coupling between EMT and BMP signalling in which EMT initiation directly primes BMP pathway competence through *SNAI2* -mediated *SMAD1* upregulation. BMP signalling, when activated, then coordinates cell fate proportions and invasion dynamics to drive collective cell invasion during vertebrate development.

## Results

### *Ex vivo* induction of EMT in chick tissue explants

We first asked whether plating tissue explants onto fibronectin-coated glass is sufficient to drive EMT and cell migration. To compare different initial EMT states, we compared two gastrula-stage (HH4) chick tissues. Firstly, the prospective neural plate (PNP) tissue, which is neuroectodermal and not fated to undergo EMT. Secondly, the anterior primitive streak (APS) tissue, which is mesodermal and already actively undergoing EMT in the HH4-stage embryo. Both PNP and APS explants were plated on human fibronectin-coated glass-bottomed dishes and incubated at 37 °C and 5.5% CO_2_ for 24 hours (Fig. 1A). Live imaging revealed robust migration in both PNP and APS explants, with cells collectively dispersing radially from the explant edge and moving over the fibronectin substrate (Fig. 1B; Supplementary Movie 1). This experimental system therefore provides a platform to determine the transcriptional events downstream of EMT initiation independently of other developmental cues.

**Fig. 1.**
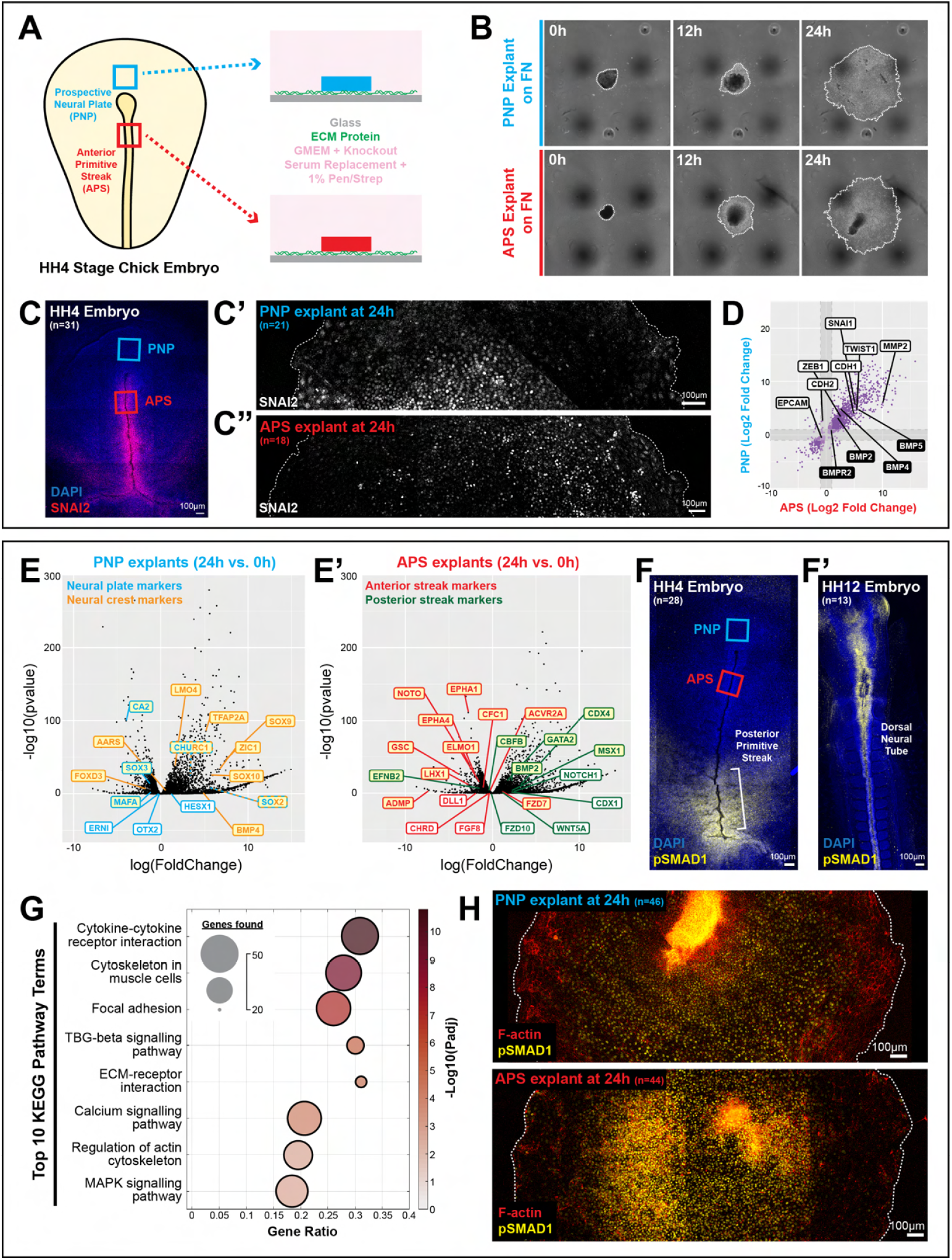
*Ex vivo* EMT assay reveals BMP activation and BMP-induced cell fate changes. A) Schematic of *ex vivo* EMT assay in chick explants. B) Live-imaging of migrating explants on fibronectin. C-C”) Representative SNAI2 immunofluorescence in HH4 embryo (C), PNP explant after 24h (C’), and APS explant after 24h (C”). D) Bulk RNA-seq fold change plot of EMT genes (white) and BMP signalling genes (black). Log2 of gene expression fold change in 24h migrating explants compared to 0h controls in APS tissue (x axis) and PNP tissue (y axis). All p-adj<0.01. Shaded: |fold change|<1. n=16-27 explants, 3 biological replicates. E-E’) Volcano plots of differentially expressed genes in PNP (E) and APS (E’) migrating explants at 24h vs 0h. Yellow highlight: p-adj<0.01. F-F’) Representative phospho-SMAD1 immunofluorescence in HH4 (F) and HH12 (F’) embryos. G) Top 8 terms enriched in gene ontology analysis of genes upregulated ≥ 2-fold in both PNP and APS explants upon migration. KEGG: Kyoto Encyclopedia of Genes and Genomes. H) Representative phospho-SMAD1 immunofluorescence and phalloidin staining in PNP (top) and APS (bottom) migrating explants after 24h.

In normal chick development, SNAI2, a key EMT driver known to be upstream of the EMT programme in both the chick mesoderm and neural crest (Nieto et al., 1994), is expressed in the primitive streak (Fig. 1C). Both PNP and APS migrating explants at 24 hours of culture showed nuclear SNAI2 protein expression, indicating activation of the EMT transcriptional programme in PNP explants, in addition to maintained expression in APS explants (Fig. 1C’,C”). To further validate EMT induction, we examined epithelial polarity in PNP migrating explants. *In vivo*, the neuroepithelial PNP region displays classical apical-basal polarity with Zonula Occludens-1 (ZO-1) and atypical Protein Kinase C (aPKC) localized to the apical domains (Fig. S1). After 24 hours in culture, however, PNP migrating explants exhibited disorganized ZO-1 and aPKC localization along the apical and basolateral domains (Fig. S1), consistent with loss of epithelial apical-basal polarity during EMT transitions. Bulk RNA sequencing performed on PNP and APS explants at 0 and 24 hours of culture (PNP-0h, PNP-24h, APS-0h, APS-24h) confirmed these morphological findings (Table S1-3). Both PNP and APS explants at 24 hours of culture showed upregulation of a common set of key EMT drivers (*SNAI1, ZEB1, TWIST1*) and a matrix metalloproteinase gene known to be implicated in EMT (Duong and Erickson, 2004), *MMP2* (Fig. 1D). These transcriptomic signatures confirmed the progression of EMT in both tissue types, demonstrating that this substrate-induced perturbation triggers a shared EMT programme across distinct germ layers. This experimental system therefore provides a robust platform to examine the immediate consequences of EMT induction.

### *Ex vivo* induction of EMT drives convergent cell fate shifts toward migratory lineages

Having established that our *ex vivo* system successfully induces EMT, we next explored the common transcriptional signatures associated with an induced migratory state. Global analysis of the transcriptomes revealed several key insights. A heatmap of the top differentially expressed genes across all four conditions showed consistent patterns, with the translation elongation gene *EEF1A1* expression enriched in 0-hour samples and the actin cytoskeletal gene *ACTB* expression in 24-hour samples, regardless of tissue type (Fig. S2A). Notably, most genes showed coordinated changes regardless of tissue type; genes upregulated in APS migrating explants were likely also upregulated in PNP explants, and vice versa (Fig. 1D). This strong correlation suggested a shared migrating explant programme of gene expression changes. Principal component analysis confirmed this pattern, with the major axis of variation (PC1) segregating samples by timepoint (Fig. S2B). Gene ontology analysis revealed that the top 500 genes contributing to PC1 were enriched for biosynthetic processes such as replication, transcription, and translation (Fig. S2B’). Meanwhile, PC2 separated samples by tissue type with enrichment of differentiation-related terms (Fig. SB,B”). Notably, 24-hour migrating samples showed greater divergence along PC2 than 0-hour samples (Fig. S2B), suggesting that while EMT activation drives a shared programme, distinct lineage identities continued to specify and diverge.

Critically, both tissues showed cell fate specification responses following EMT activation, with shifts toward more migratory fates within their respective germ layers. PNP explants downregulated neural plate markers and upregulated neural crest markers after 24 hours in culture (Fig. 1E), suggesting a fate shift toward the migratory neural crest lineage. In parallel, APS explants showed downregulation of anterior primitive streak markers and upregulation of posterior streak markers (Fig. 1E’). Together, these findings demonstrate that substrate-induced EMT activation triggers cell fate responses across germ layers.

### *Ex vivo* induction of EMT triggers BMP pathway activation

We noticed that both neural crest and posterior primitive streak represent migratory cell fates within their respective germ layers, and both are known to be promoted by BMP signalling during normal development (Marchant et al., 1998; Mayor et al., 1995; Wilson et al., 1997; Neave et al., 1997; Nguyen et al., 1998; Glavic et al., 2004; Liem et al., 1995; Selleck et al., 1998; James and Schultheiss, 2005; Faure et al., 2002; Dale et al., 1992; CM et al., 1992) (Fig. 1F,F’). To explore this further, we first examined whether EMT induction affected the expression of BMP pathway components. Comparison of shared patterns of gene expression changes in both PNP and APS explants after 24 hours revealed up-regulation of BMP ligand genes (*BMP2, BMP4, BMP5*) (Fig. 1D & S3B). Additionally, many ECM components and remodeling enzymes were upregulated (Fig. S3A), consistent with the migratory phenotype. Gene ontology analysis of genes upregulated at least 2-fold in both tissues confirmed enrichment for the TGF*β* signalling pathway (Fig. 1G & S4), which includes the BMP signalling pathway. To directly assess BMP signalling activity, we performed immunostaining for phosphorylated SMAD1 (pSMAD1), a well-established readout of active BMP signalling (Kretzschmar et al., 1997). Both PNP and APS migrating explants showed strong nuclear pSMAD1 signal (Fig. 1H), confirming pathway activation following EMT induction. This BMP signalling activation in PNP and APS migrating explants was observed regardless of the ECM substrate used, occurring on both human fibronectin and rat-tail collagen I (Fig. S5). Together, these results demonstrate that EMT initiation triggers both BMP pathway component expression and signalling activity, consistent with the observed cell fate changes toward migratory lineages.

### *SMAD1* upregulation couples EMT initiation to BMP signalling competence

The observed BMP pathway activation following EMT induction raised a key question: are these processes mechanistically linked, or do they operate in parallel? To investigate this, we searched our bulk RNA sequencing dataset for genes whose expression might control BMP signalling competence. We reasoned that genes linking EMT to BMP signalling should be upregulated only in PNP migrating explants, which undergo EMT induction, and not in APS explants, which maintain pre-existing EMT programmes. *SMAD1* mRNA was significantly upregulated in PNP explants after 24 hours of migration compared to 0 hours, consistent with BMP pathway activation following EMT induction (Fig. 2A). SMAD1 protein functions as an intracellular mediator of BMP signalling, without which cells cannot transduce BMP ligand signals to activate downstream target gene expression. The regulation of *SMAD1* expression therefore represents a mechanism for increasing cellular BMP competence during EMT.

**Fig. 2.**
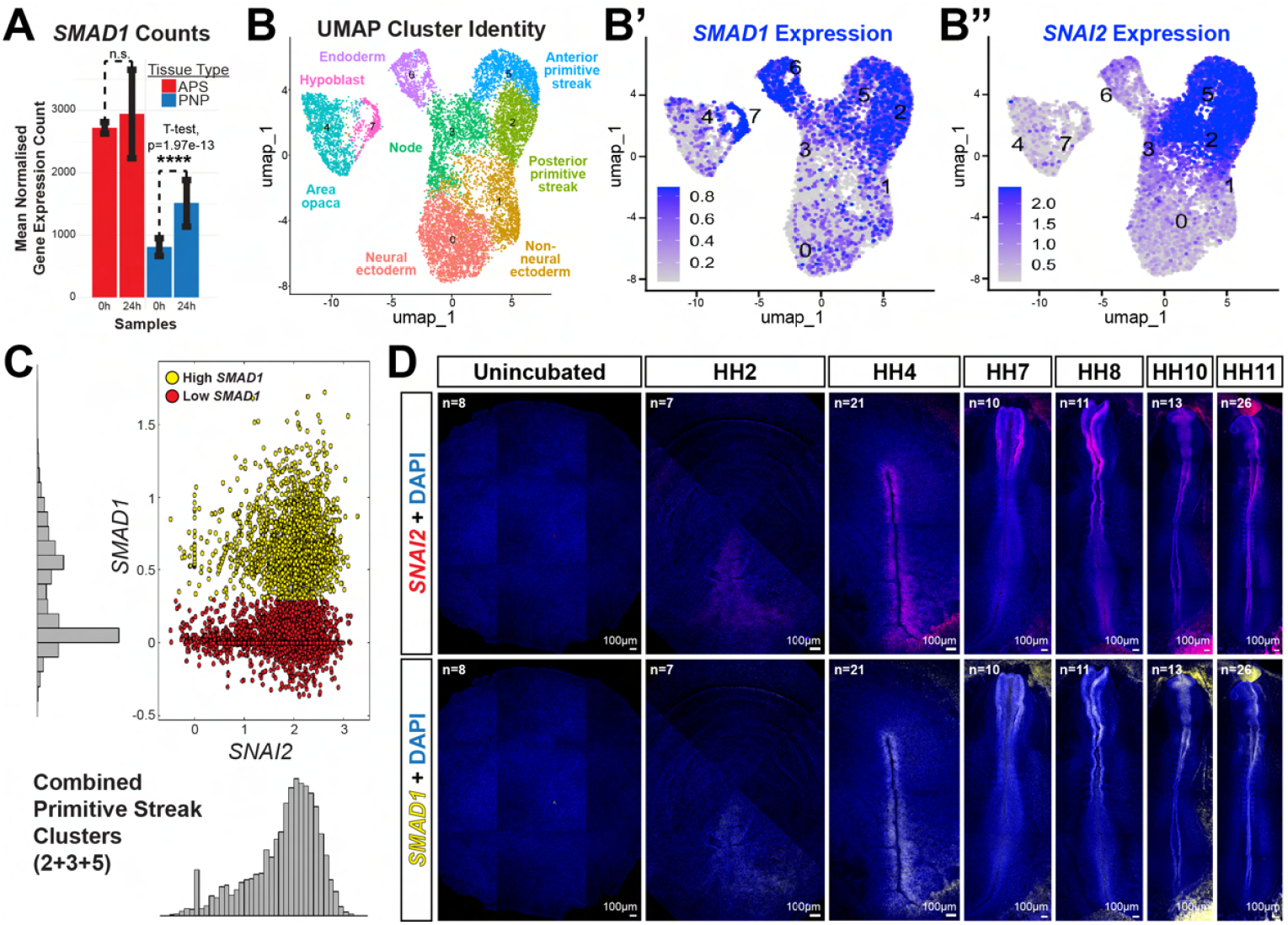
*SNAI2* and *SMAD1* expression correlates *in vivo*. A) Bulk RNA-seq *SMAD1* expression in PNP and APS explants (0h vs 24h). B-B”) scRNA-seq of gastrula-stage chick embryos. UMAP clusters (B), *SMAD1* expression (B’), *SNAI2* expression (B”). Sex and cell cycle regressed. Dim 1:11, res 0.18. C) Distrbution of *SMAD1* and *SNAI2* expression in a combined primitive streak cluster (clusters 2+3+5). Color shows the bimodal distribution of *SMAD1* expression. Red: Cells with high *SNAI2* and low *SMAD1*. Yellow: Cells with high *SNAI2* and high *SMAD1*. D) Representative *SNAI2* and *SMAD1 in situ* HCR in whole embryos. Bleed-through was tested and controlled for by changing HCR hairpins.

Interestingly, *SMAD1* expression remained constitutively high in APS explants at both timepoints, suggesting that this tissue already possessed the molecular machinery for BMP pathway competence prior to explantation and migration (Fig. 2A). As APS explants also upregulate BMP signalling in migrating explants, we asked whether this activation resulted from removing tissues from embryonic sources of BMP inhibition. During early chick development, BMP inhibitors such as *CHRD* are expressed and secreted from the Hensen’s node (Fig. S6A) (Streit et al., 1998), raising the possibility that dissection alone might be sufficient to relieve BMP inhibition. To test this, we cultured explants in suspension for 24 hours, thereby removing both ECM contact and the physical substrate required for migration while still isolating tissues from embryonic sources of BMP inhibitors. Floating APS explants, which already express EMT factors *in vivo*, gained BMP signalling activity after 24 hours (Fig. S6B,B’), consistent with relief from node-derived inhibition. In contrast, floating PNP explants lacked detectable pSMAD1 signal (Fig. S6B,B’), indicating that BMP activation in PNP explants requires more than the removal of exposure to endogenous BMP inhibitors. We reason that this is likely due to the induced *SMAD1* expression in PNP explants, suggesting that *SMAD1* transcription could be act as a molecular link between EMT initiation and BMP signalling competence.

### Identification of *SNAI2* as a candidate regulator of *SMAD1* expression

To determine candidates for the control of *SMAD1* expression *in vivo*, we generated single-cell RNA sequencing data from gastrula-stage chick embryos. Pearson’s correlation analysis revealed the top genes positively correlated with *SMAD1* expression (Table S4). Strikingly, among the highly correlated genes, *SNAI2* emerged as a compelling candidate, ranking 20th with a Pearson’s correlation coefficient of 0.26. Notably, *SNAI2* was the only well-established EMT transcription factor ranked highly on this list; other core EMT drivers showed substantially weaker correlations with *SMAD1* (*ZEB1* : rank 104, r=0.14; *TWIST1* : rank 149, r=0.11; *ZEB2* : rank 278, r=0.07; *SNAI1* : rank 1261, r=0.01). This correlation-based finding identified *SNAI2*, a well-characterized EMT driver in both chick mesoderm and neural crest (Nieto et al., 1994), as a data-driven candidate for upstream regulation of *SMAD1* expression during EMT initiation.

UMAP clustering identified distinct embryonic territories (Fig. 2B & S7; Table S5-12), with overlapping expression of *SMAD1* and *SNAI2* transcripts specifically within the primitive streak clusters which correspond to precisely the embryonic regions of EMT initiation (Fig. 2B’,B”). Detailed correlation analysis within a combined primitive streak cluster (clusters 2+3+5) revealed two distinct cell populations: one expressing high *SNAI2* but low *SMAD1*, and another coexpressing both *SNAI2* and *SMAD1* at high levels (Fig. 2C). This bimodal distribution of *SMAD1* expression in *SNAI2* -expressing cells supports a sequential activation model in which *SNAI2* expression precedes and potentially drives *SMAD1* expression in a subset of cells undergoing EMT initiation. A third cell population expressing low *SNAI2* and high *SMAD1* was observed in some other clusters (Fig. S8). To validate these findings, we performed multiplexed *in situ* hybridization chain reaction (HCR) for *SNAI2* and *SMAD1* across chick embryonic development. These experiments confirmed spatiotemporal co-localization of *SNAI2* and *SMAD1* transcripts in the primitive streak, neural plate border, neural folds, and dorsal neural tube regions (Fig. 2D) across development, consistent with literature (Gont and Lough, 2000; Le Dréau et al., 2012; Acloque et al., 2011; Hardy et al., 2011; McLarren et al., 2003). This establishes that *SNAI2* and *SMAD1* coexpression is a consistent feature of embryonic regions of EMT initiation. Together, these findings demonstrate that *SNAI2* and *SMAD1* are co-expressed during development in a pattern consistent with a transcriptional hierarchy, where *SNAI2* expression precedes *SMAD1* upregulation within regions of EMT initiation.

### BMP signalling activity is restricted to regions of EMT initiation during development

Having established that *SNAI2* and *SMAD1* transcripts co-localize at EMT initiation sites, we next asked whether BMP signalling activity itself shows similar spatial restriction. Interestingly, examination of pSMAD1 signal distribution in *ex vivo* PNP and APS migrating explants revealed that nuclear pSMAD1 signal was strongest in the center of the explants and progressively reduced toward the periphery where cells displayed more mesenchymal morphology (Fig. S9). This gradient pattern suggested that BMP signalling activity might be spatially restricted to regions of active EMT initiation rather than maintained throughout migration. To investigate this, we compared the spatial dynamics of SNAI2 and pSMAD1 expression across chick embryonic development from gastrulation through neural crest delamination. Immunostaining of SNAI2 in HH4 to HH11 chick embryos revealed nuclear signal in the primitive streak, migrating mesoderm, neural folds, dorsal neural tube, and migrating neural crest cells (Fig. 3A). In parallel, pSMAD1 immunostaining revealed nuclear signal in the posterior primitive streak, neural folds, and dorsal neural tube (Fig. 3A). Cryosectioning and confocal reslices revealed that SNAI2 signal initiates in the dorsal neural tube and persists in delaminated and migrating neural crest cells (Fig. 3B & S10). In contrast, pSMAD1 signal was sharply restricted to the dorsal neural tube itself, with little to no signal detectable in the delaminated neural crest cells that had already begun migration (Fig. 3B & S10). Similarly, SNAI2 protein expression initiated in the primitive streak and persisted in migrating mesoderm cells (Fig. 3C), but pSMAD1 signal was restricted to the epiblast and primitive streak regions and was absent from migrating mesoderm cells (Fig. 3C). These observations confirm that EMT initiation and BMP signalling activity are tightly coupled *in vivo*, with BMP signalling restricted to regions of EMT initiation during normal development (Fig. 3D).

**Fig. 3.**
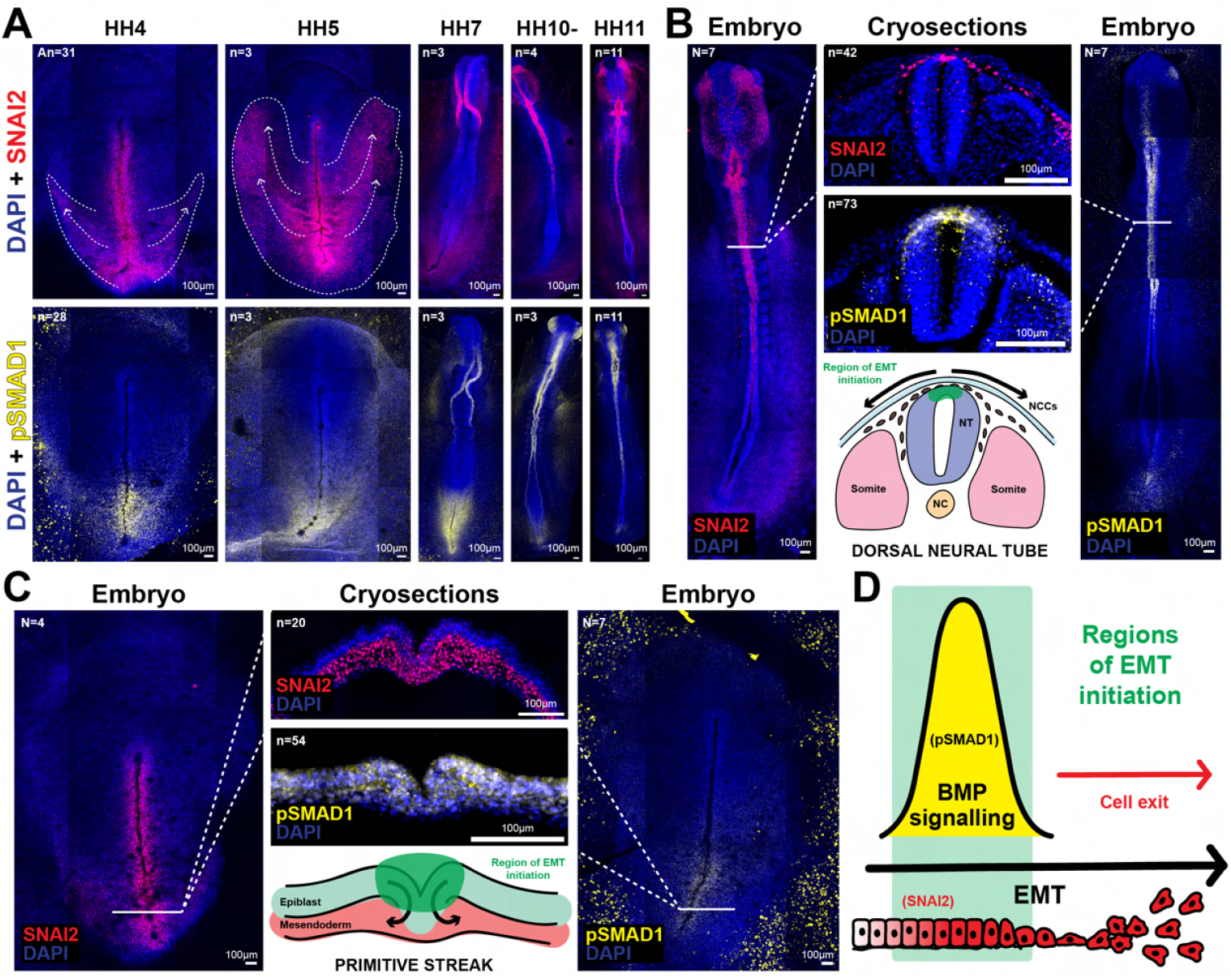
BMP signalling is restricted to regions of EMT initiation *in vivo*. A) Representative SNAI2 and pSMAD1 immunofluorescence in whole embryos. Embryos were imaged from the epiblast side, but SNAI2 immunofluorescence signal from the migrating mesoderm layer can be seen, as indicated by the dashed white lines and arrows. B) Representative SNAI2 and pSMAD1 immunofluorescence in HH10 embryos and cryosections during neural crest delamination. C) Representative SNAI2 and pSMAD1 immunofluorescence in HH4 embryos and cryosections during mesoderm ingression. D) Schematic of BMP signalling patterns coordinated with EMT progression.

### *SNAI2* expression is sufficient to drive *SMAD1* expression *in vivo*

We next asked whether *SNAI2*, as a transcription factor, might regulate *SMAD1* transcription. Although the Snail family of transcription factors canonically function as transcriptional repressors (reviewed in Nieto, 2002), there have been reports of these transcription factors acting as transcriptional activators (reviewed in de Herreros, 2024). To test whether *SNAI2* expression is sufficient to induce *SMAD1* expression, we performed targeted electroporation of a *SNAI2* -overexpression construct with nuclear EGFP reporter (pCIG-*SNAI2*) into the PNP region of HH4-stage chick embryos, a region that normally expresses neither *SNAI2* nor *SMAD1*. As a negative control, embryos were electroporated with the pCIG backbone alone, which expresses nuclear EGFP but lacks the *SNAI2* coding sequence (Fig. 4A). Constructs were coinjected with brilliant blue dye into the space between the epiblast and the vitelline membrane to visualize successful targeting (Fig. S11A). After 8 hours of incubation, we validated ectopic SNAI2 expression in EGFP-positive cells using SNAI2 immunofluorescence. While control embryos showed no detectable SNAI2 protein in the electroporated PNP region, embryos electroporated with pCIG-*SNAI2* exhibited robust nuclear SNAI2 protein expression specifically in EGFP-expressing cells (Fig. 4B), confirming successful overexpression and nuclear localization of the transcription factor. We further validated ectopic *SNAI2* mRNA expression using HCR staining (Fig. 4C). Imaging revealed that *SNAI2* -overexpressing cells remained within the prospective neural plate without undergoing ingression, consistent with previous reports (Moly et al., 2016). We then performed multiplexed *in situ* HCR for both *SNAI2* and *SMAD1* transcripts. As a positive control of *SMAD1* expression, endoderm cells (naturally *SNAI2* -negative and *SMAD1* -positive) were stained (Fig. S11B,C). As a negative control, cells electroporated with the pCIG backbone were stained and showed no expression of neither *SNAI2* nor *SMAD1* in EGFP-positive cells within the electroporated PNP region (Fig. 4D,E & S11C). In contrast, EGFP-positive cells electroporated with the pCIG-*SNAI2* construct co-expressed both *SNAI2* and *SMAD1* (Fig. 4D,E & S11C), demonstrating that ectopic *SNAI2* expression was sufficient to drive ectopic expression of *SMAD1*. Any potential fluorophore bleed-through was tested and controlled for by changing HCR hairpins (Fig. S11C).

**Fig. 4.**
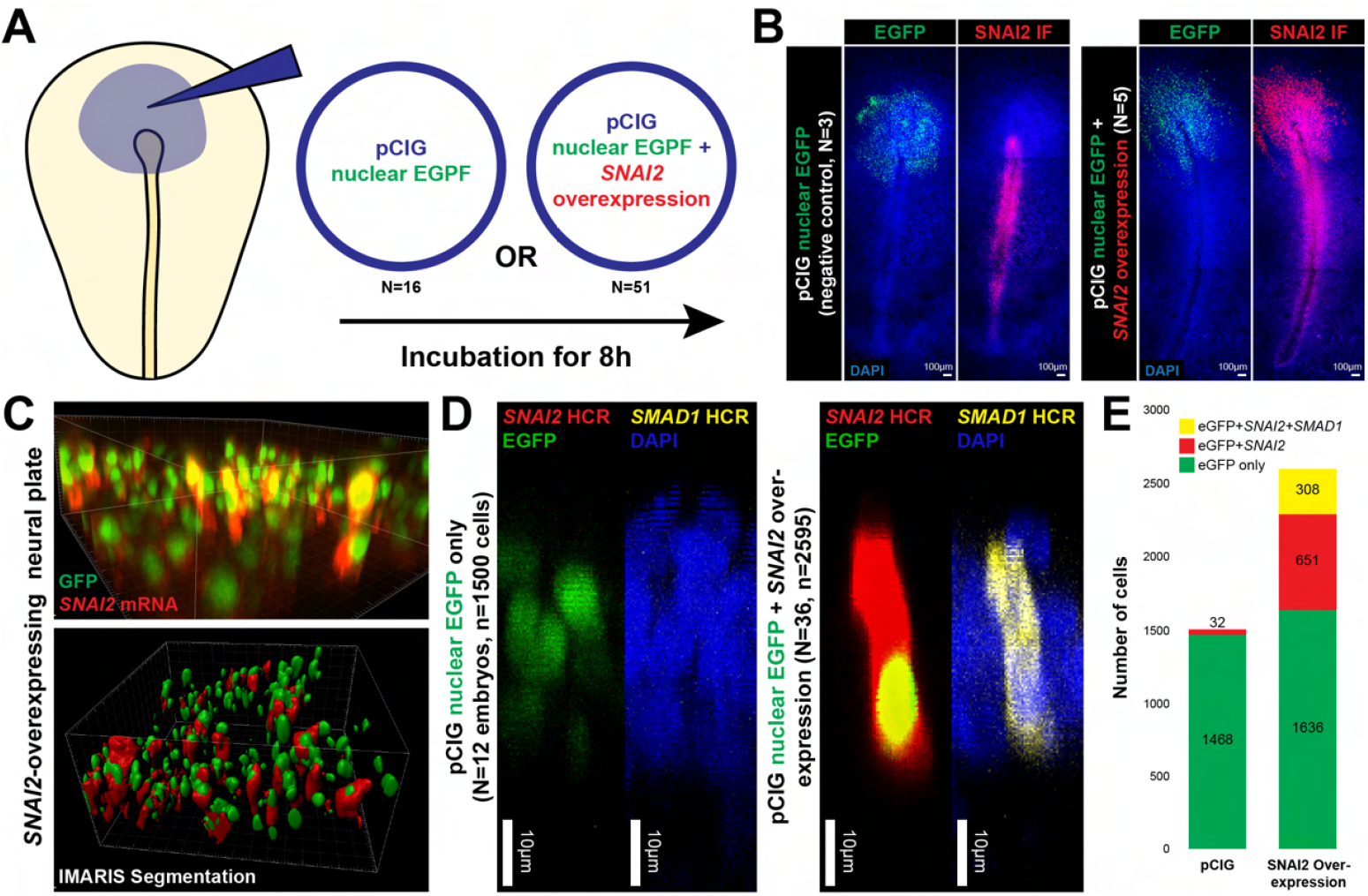
*SNAI2* overexpression sufficient to drive *SMAD1* expression *in vivo*. A) Schematic of *SNAI2* overexpression experiment in HH4 prospective neural plate. Embryos were electroporated with pCIG (negative control) or pCIG-*SNAI2* (*SNAI2* overexpression construct) and incubated for 8h. B) Representative SNAI2 immunofluorescence in pCIG vs pCIG-*SNAI2* electroporated embryos. C) IMARIS 3D view snapshots of *SNAI2* -overexpressing prospective neural plate cells. D) FIJI 3D view of representative *SNAI2* and *SMAD1 in situ* HCR in pCIG vs pCIG-*SNAI2* electroporated cells. E) Cell counts. The *SNAI2* -positive cells in the pCIG controls are due to some node cells being electroporated.

To test if *SNAI2* is required for *SMAD1* expression, we used CRISPR-Cas9-mediated gene editing to knock out *SNAI2* expression in regions where it is normally expressed. We cloned *SNAI2* single guide RNAs (sgRNA) into the pcU6-3 plasmid and coelectroporated these with the pCAG Cas9-2A-Citrine plasmid into the neural plate border region of HH4-stage embryos (Fig. S12C). The sgRNA was designed to target a DpnII restriction enzyme cut site within the *SNAI2* gene, such that successful CRISPR-Cas9 editing would disrupt these sites and prevent DpnII digestion (Fig. S12A). PCR amplification of the target sequence followed by DpnII digestion and gel electrophoresis confirmed effective gene editing (Fig. S12B). As a negative control, embryos were electroporated with the pCAG Cas9-2A-Citrine vector alone (Fig. S12C). Plasmids were co-electroporated with brilliant blue dye to ensure successful targeting (Fig. S12D), and embryos were incubated for 24 hours to allow sufficient time for gene editing and subsequent transcriptional changes. Since only the pCAG Cas9-2A-Citrine vector contains a fluorescent reporter (Citrine), *SNAI2* knockout is necessarily mosaic even within Citrine-positive cells, as we cannot definitively identify which cells received both the pCAG Cas9-2A-Citrine plasmid and the pcU6-3 *SNAI2* -sgRNA plasmid. Despite this inherent variability, quantitative analysis using cell surface segmentation revealed measurable changes in the targeted cell populations. CRISPR-mutagenized embryos showed a significant decrease in SNAI2 protein levels in Citrine-positive cells compared to controls (Fig. S12E), confirming effective knockdown of SNAI2 expression. Similarly, *SNAI2* transcript levels were significantly reduced in the CRISPR-mutagenized embryos compared to controls (Fig. S12G), validating the gene editing approach at both protein and mRNA levels. However, despite this reduction in *SNAI2* expression, *SNAI2* knockout samples showed no significant changes in either pSMAD1 protein levels (Fig. S12F) or *SMAD1* transcript levels (Fig. S12H) compared to controls. Together, these results demonstrate that *SNAI2* overexpression is sufficient to drive *SMAD1* expression *in vivo*. However, the observation that some cells express *SMAD1* without *SNAI2* in our single-cell RNA sequencing data (Fig. 2B’,B” and Fig. S8) suggests that other transcriptional regulators or signalling pathways may also regulate *SMAD1* expression.

### BMP signalling is dispensable for EMT initiation but regulates cell fate proportions

Having established that EMT activation induces BMP pathway competence and that *SNAI2* can drive *SMAD1* expression, we next investigated the functional consequences of this molecular coupling. We first treated whole HH4-stage chick embryos with the pan BMP type I receptor (ALK1/2/3/6) inhibitor LDN193189 (LDN), at 100*µ*M in EC culture agaralbumen) and let the embryos grow overnight. As expected, BMP inhibition caused extensive neural tube closure defects, with embryos displaying persistently open neural plates (Fig. 5A). Quantification revealed two prominent LDN-induced phenotypes: a significant increase in lateral somite columns beyond the normal bilateral pair (Fig. 5B) and failure of neural fold convergence, with significantly increased distance between neural folds (Fig. 5B’). Cryosections confirmed the formation of additional somites lateral to the main body axis (Fig. 5A). These morphological defects demonstrate that BMP signalling is essential for coordinating the complex morphogenetic movements and patterning events that shape normal embryonic development.

**Fig. 5.**
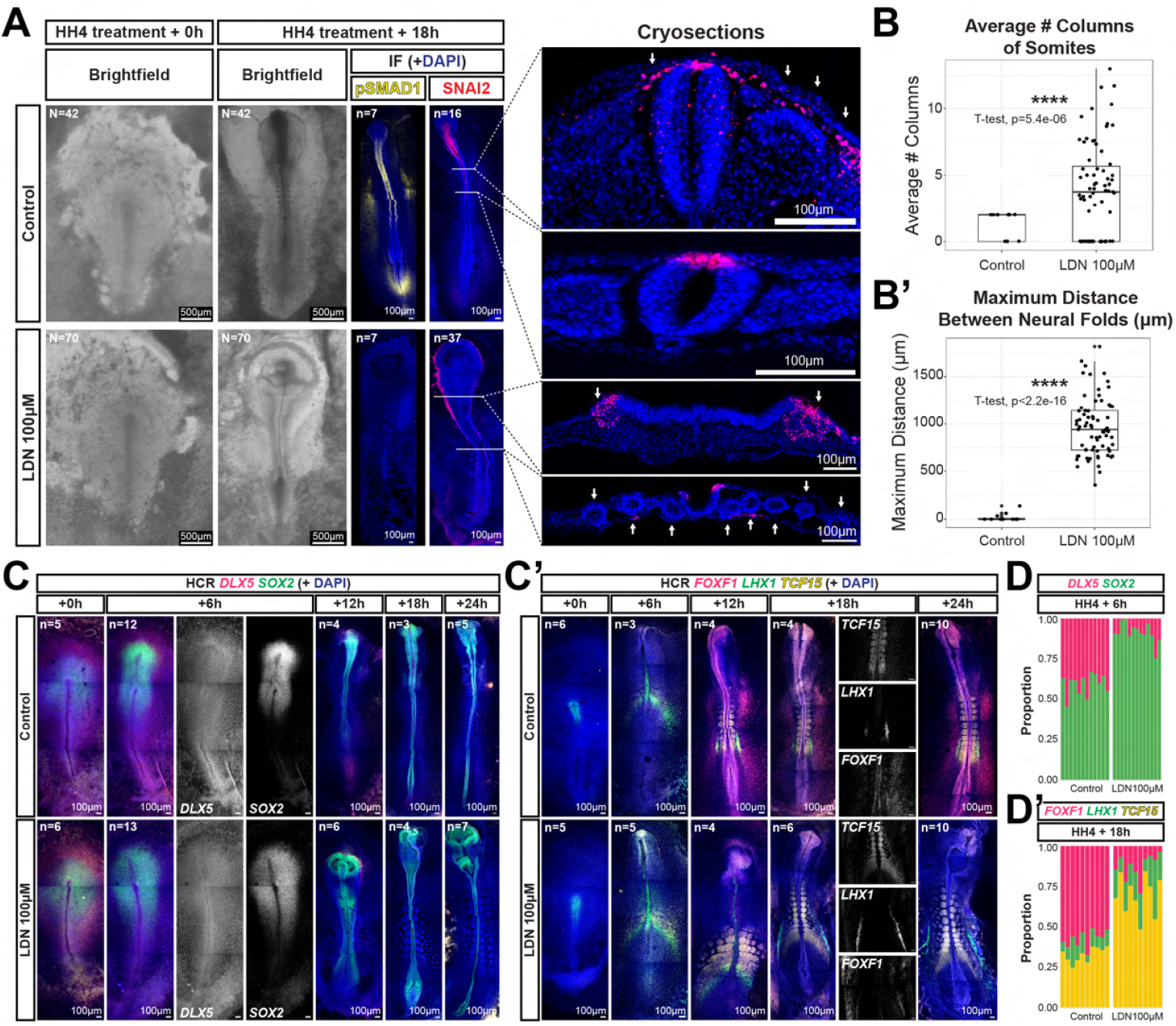
BMP signalling tunes cell fate proportions but not EMT initiation. A) Representative brightfield and SNAI2/pSMAD1 immunofluorescence in control and LDN-treated embryos (left). Representative cryosections of SNAI2 immunofluorescence in control and LDN-treated embryos (right). In the top cryosection panel, arrows point to SNAI2-positive migrating neural crest cells. In the second-to-bottom cryosection panel, arrows point to SNAI2-positive cells migrating from the flat neural plate border region. In the bottom panel, arrows point to extra lateral somites. B-B’) Quantification of somite columns (B) and neural fold distance (B’) in control vs LDN-treated embryos. For somite columns, WT should be 2 (1 column of somites on each side of the main body axis), unless development is delayed. For neural fold distance, WT should be 0 *µ*m (successful neural tube closure), unless development is delayed. C-C’) neural plate border marker *DLX5* + neural plate marker *SOX2* (C) and lateral plate mesoderm marker *FOXF1* + intermediate mesoderm marker *LHX1* + paraxial mesoderm marker *TCF15* (C’) *in situ* HCR in control vs LDN-treated embryos at 0h, 6h, 12h, 18h, and 24h of incubation. D-D’) Quantification of proportional embryonic area positive for *DLX5* vs *SOX2* at 6h (D) and *FOXF1* vs *LHX1* vs *TCF15* at 18h (D’) in control vs LDN-treated embryos. Proportions were calculated relative to the total combined area of all measured domains (neural plate border + neural plate for D; paraxial + intermediate + lateral plate mesoderm for D’).

Critically, LDN-treated embryos maintained robust SNAI2 protein expression in the neural plate border region despite complete loss of pSMAD1 signal (Fig. 5A), confirming that BMP signalling is dispensable for EMT initiation. Cryosections confirmed extensive migration of SNAI2-positive cells from the flattened neural plate border region (Fig. 5A), demonstrating that EMT can proceed even in the absence of BMP signalling and proper neural tube morphogenesis. Moreover, multiplexed *in situ* HCR revealed that LDN-treated embryos expressed *SNAI2, SMAD1*, dorsal neural tube marker *BMP4*, pre-migratory neural crest marker *FOXD3*, and migratory neural crest marker *SOX10*, all within the neural plate border region (Fig. S13A). This expression pattern persisted even when LDN treatment was performed earlier on HH3/3+ stage embryos (Fig. S13B), demonstrating that the EMT transcriptional programme can be activated independently of BMP signalling across multiple developmental timepoints. These findings are mirrored *in vitro*, as 10nM LDN treatment did not prevent the initiation of EMT in either PNP or APS tissue (Fig. S14). This suggests that BMP signalling is not required for the onset of migration, but may instead function to regulate other aspects of cellular behaviour during cell invasion.

However, while EMT initiation proceeded normally, BMP inhibition significantly altered cell fate proportions in the embryo. By 6 hours post-treatment, LDN-treated embryos showed reduced expression of the neural plate border marker *DLX5* and expanded expression of the neural plate marker *SOX2* (Fig. 5C,D), indicating a shift toward neural plate identity at the expense of neural plate border fate. Similarly, analysis of mesodermal markers revealed that BMP inhibition reduced the expression of lateral plate mesoderm marker *FOXF1* while expanding the expression of intermediate mesoderm marker *LHX1* and paraxial mesoderm marker *TCF15* by 18 hours post-treatment (Fig. 5C’,D’). These cell fate shifts are consistent with known roles of BMP signalling in promoting lateral plate mesoderm and neural plate border specification during normal development. Together, these results demonstrate that BMP signalling is dispensable for EMT initiation but essential for regulating cell fate proportions, specifically promoting migratory lateral identities (neural plate border and lateral plate mesoderm) over more medial identities (neural plate and paraxial mesoderm).

### BMP signalling modulates cell invasion dynamics

Having established that BMP signalling is dispensable for EMT initiation, we next asked whether BMP signalling regulates collective cell invasion downstream of EMT. To this end, we compared tissues already undergoing EMT with different endogenous BMP signalling states. Specifically, we compared APS tissue, which lacks BMP signalling, with the PPS tissue, which is BMP-positive. We first treated BMP-negative APS explants with recombinant mouse BMP4 protein (100 ng/ml) (Fig. 6A). Live imaging revealed that BMP4 treatment significantly enhanced APS explant spreading, with treated explants displaying faster initial dispersal, greater final spread area, and more complete spreading compared to controls (Fig. 6B & S15; Supplementary Movie 2). Conversely, LDN treatment of BMP-positive PPS explants had reduced spreading, with inhibited explants showing slower dispersal dynamics and less complete spreading (Fig. 6B & S15; Supplementary Movie 2). These results demonstrate that BMP signalling directly modulates cell dispersal behaviour in tissues already undergoing EMT, supporting a downstream rather than initiating role for BMP signalling.

**Fig. 6.**
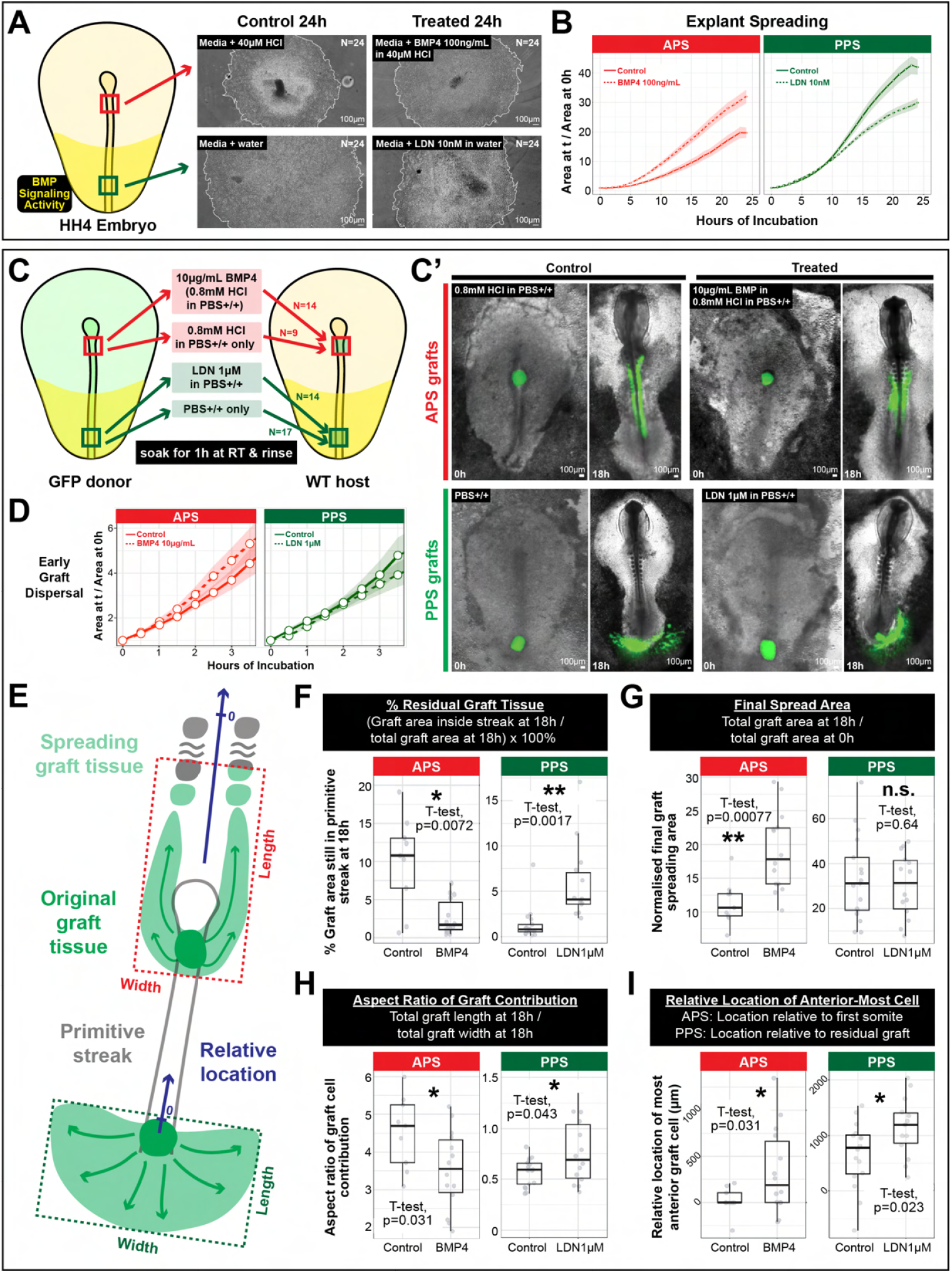
BMP signalling coordinates invasion dynamics. A) Explant experiment schematic, representative images of control and treated APS and PPS explants. B) Area quantification over time. C-C’) Grafting experiment schematic (C) and representative snapshots of grafted embryos at 0h and 18h of incubation (C’). D) Area quantification of early graft dispersal. E) Schematic of different types of quantification performed on APS and PPS grafts. F) Quantification of the percent of graft tissue remaining inside the primitive streak at end timepoint. G) Quantification of the final graft spreading area, normalised by the area at 0h. H) Quantification of the graft aspect ratio (length divided by width) at the end timepoint. I) Quantification of the degree of graft anterior migration, relative to the location of the first somite for APS grafts and to the location of the original graft location for PPS grafts.

To validate these findings in an embryonic context, we performed *in vivo* grafting experiments. APS and PPS tissue were dissected from GFP donor embryos and soaked in either BMP4 (10 *µ*g/mL) or LDN (1 *µ*M) respectively for 1h at RT, then grafted into WT host embryos and monitored by live imaging (Fig. 6C,C’). Consistent with our *ex vivo* results, BMP4-treated APS grafts showed accelerated dispersal during the first few hours of incubation (Fig. 6D). Furthermore, analysis of graft tissue retention within the primitive streak revealed that BMP signalling modulates the rate of precursor region depletion. BMP4-treated APS grafts showed significantly reduced retention of graft tissue within the primitive streak after 18h of incubation (Fig. 6E,F), indicating faster depletion of the precursor region. However, the final spread area was similar between BMP4-treated and control APS grafts after 18h of incubation (Fig. 6E,G). Analysis of graft contribution patterns also revealed that BMP signalling also influences migration directionality. BMP4-treated APS grafts showed decreased aspect ratio (length/width) compared to controls (Fig. 6E,H), with significantly increased anterior migration (Fig. S16) and increased lateral migration (Fig. S16). This enhanced dispersal in both anterior and lateral directions is consistent with specification of lateral mesoderm fates as well as faster exit from the primitive streak.

Conversely, LDN-treated PPS grafts showed initially slower cell dispersal during the first few hours of incubation (Fig. 6D). Analysis of PPS graft retention revealed that LDN treatment led to increased tissue retention in the primitive streak compared to controls (Fig. 6E,F), indicating slower precursor region depletion. The final spread area was similar between LDN-treated and control PPS grafts after 18h of incubation (Fig. 6E,G). LDN-treated PPS grafts showed decreased aspect ratio compared to controls (Fig. 6E,H), with decreased anterior migration (Fig. 6E,I), consistent with the shift toward more embryonic mesodermal fates that contribute along a narrower anterior-posterior axis. Together, these *in vivo* grafting experiments demonstrate that BMP signalling modulates multiple aspects of collective cell invasion in tissues already undergoing EMT: it enhances cell dispersal rates, regulates precursor region depletion rates, and influences migration directionality.

All together, our findings reveal a reciprocal coupling between EMT initiation and BMP signalling that coordinates collective cell invasion during vertebrate development. EMT initiation directly primes BMP pathway competence through *SNAI2* -mediated *SMAD1* up-regulation. When activated, BMP signalling in turn regulates cell fate proportions and invasion dynamics—including cell dispersal dynamics, precursor region depletion rates, and migration directionality. However, our findings are also consistent with known roles for BMP signalling in regulating cell fate specification, typically understood to be upstream of EMT initiation. This coupling provides a molecular mechanism for synchronising fate specification with invasion progression, positioning EMT as a regulatory process that increases BMP competence to coordinate tissue-level behaviours during collective cell invasion.

## Discussion

### EMT initiation triggers BMP pathway competence to coordinate collective cell invasion

The traditional view of EMT positions morphogen signalling upstream of a linear cascade: morphogens first specify cell fates, which then trigger EMT programs, leading to cell invasion (reviewed in Acloque et al., 2009; Thiery et al., 2009; Leathers and Rogers, 2022; Zhang et al., 2025). Our results add to this model by demonstrating that EMT initiation itself can directly prime BMP pathway competence, creating reciprocal interactions between EMT and morphogen competence that coordinate collective cell invasion at the population level. Therefore, our findings reveal reciprocal coupling between EMT and BMP-mediated cell fate specification during collective cell invasion.

By forcing cell migration *ex vivo*, we demonstrate that EMT initiation (marked by SNAI2 expression) triggers BMP pathway activation. This coupling operates in both neuroectodermal and mesodermal tissues, with substrate-induced EMT driving convergent cell fate changes toward migratory lineages in each germ layer: neural crest from prospective neural plate and posterior primitive streak from anterior primitive streak. The observation that these fate changes occur in both tissues despite their distinct developmental origins strongly suggests that EMT initiation provides a general mechanism for priming BMP signalling. Critically, our floating explant experiments demonstrate that this priming is functionally meaningful: PNP tissue requires active EMT induction to activate BMP signalling, whereas APS tissue (which already expresses EMT factors) activates BMP signalling upon simple removal from embryonic BMP inhibitors. This demonstrates that tissue-specific BMP competence, established through differential EMT states, determines whether cells can respond to changes in the morphogen environment.

Single-cell RNA sequencing reveals co-localization of *SNAI2* and *SMAD1* expression in regions of EMT initiation, with a bimodal distribution consistent with sequential activation where *SNAI2* expression precedes *SMAD1* upregulation in a subset of cells. Our gain-of-function experiments establish that ectopic *SNAI2* expression is sufficient to drive *SMAD1* transcription in naive tissue, directly linking the key EMT driver to BMP pathway competence. However, the mosaic CRISPR-Cas9 experiments achieved only modest reduction in *SNAI2* levels and were insufficient to test necessity. Furthermore, the observation that some cells express *SMAD1* without *SNAI2* in our single-cell data suggests that additional transcriptional regulators likely contribute to *SMAD1* expression during normal development.

A striking feature of this coupling is the sharp spatial and temporal restriction of BMP signalling activity to regions of EMT initiation. Both *in vivo* during neural crest delamination and mesoderm ingression and *ex vivo* in our migrating explants, pSMAD1 signal is highest in cells within the EMT initiation zone and progressively decreases in cells that have become mesenchymal. This spatial pattern reveals that BMP signalling functions as a transient coordinator during the transition from epithelial to migratory states, rather than as a sustained signal throughout migration. The restriction of BMP activity to EMT initiation zones may serve to temporally gate cell fate decisions to a specific developmental window, ensuring that fate specification occurs synchronously with EMT progression rather than persisting into later migratory phases.

Critically, BMP signalling proves dispensable for EMT initiation itself. LDN-mediated BMP inhibition does not prevent SNAI2 expression or the initiation of cell invasion in either *ex vivo* explants or *in vivo* embryos, establishing that EMT can proceed independently of BMP pathway activity. However, BMP signalling coordinates two critical processes: cell fate proportions and invasion dynamics. BMP inhibition shifts cell fates away from migratory lateral identities (neural plate border & lateral plate mesoderm) toward more medial identities (neural plate & paraxial mesoderm), consistent with the well-established role of BMP signalling in dorsoventral and mediolateral patterning. Additionally, BMP signalling modulates the tempo of cell dispersal, precursor region depletion, and migration directionality in tissues already undergoing EMT. This functional architecture, where EMT primes BMP competence through *SMAD1* upregulation and activated BMP signalling coordinates fate specification and invasion dynamics, provides a mechanism for synchronising cellular state transitions with morphogen-driven patterning during collective cell invasion.

### Coordinating precursor region dynamics during chick axis elongation

The coupling between EMT and BMP signalling provides a molecular mechanism for one of the fundamental challenges of vertebrate axis elongation: how to simultaneously maintain precursor populations while generating the differentiated, migratory cells needed to extend the body axis. During chick development, posterior regions must continuously contribute cells to the elongating body while maintaining sufficient precursor populations for continued growth. The posterior primitive streak exemplifies this challenge, functioning as a precursor region that must balance maintenance with the production of extraembryonic and lateral mesodermal derivatives. Our findings suggest that BMP signalling modulates this balance by tuning the rate at which precursor cells transition from maintenance to differentiation and dispersal.

Our grafting experiments directly demonstrate this regulatory function. BMP4 treatment of anterior primitive streak tissue (which lacks endogenous BMP signalling and normally contributes to paraxial mesodermal derivatives) accelerates both cell dispersal and precursor region depletion. Treated grafts show reduced retention in the primitive streak after 18 hours and enhanced contribution to anterior and lateral embryonic regions, effectively converting anterior streak behaviour to resemble posterior streak dynamics. Conversely, LDN treatment of posterior primitive streak tissue (which is naturally BMP-positive) slows initial dispersal and increases tissue retention in the primitive streak. This bidirectional modulation reveals that BMP signalling actively tunes the tempo of precursor region exit rather than simply permitting it to occur, functioning as a temporal regulator that sets the pace of tissue remodelling during axis elongation.

The role of BMP signalling in specifying cell fates along the neural plate border (reviewed in Thawani and Groves, 2020; Groves and LaBonne, 2014; Pla and Monsoro-Burq, 2018; Prasad et al., 2019) and within mesodermal tissues (reviewed in Kishigami and Mishina, 2005) is well established, but our findings reveal how this pathway is integrated with EMT dynamics to coordinate collective cell behaviours at the tissue level. Consistent with known roles in morphogenesis and patterning (Castranio and Mishina, 2009; Eom et al., 2011; Ybot-Gonzalez et al., 2007; Tonegawa and Takahashi, 1998; Dias et al., 2014), BMP inhibition with LDN leads to neural tube closure defects and ectopic lateral somite formation. Importantly, the persistence of SNAI2 expression and cell invasion from the neural plate border region in LDN-treated embryos demonstrates that individual cells retain the capacity to undergo EMT in the absence of BMP activity. However, without BMP-mediated coordination of fate specification and invasion dynamics, these EMT events become decoupled from tissue-scale organization: the neural plate remains open with cells dispersing from flattened folds, and mesodermal precursors produce excess paraxial mesoderm at the expense of lateral plate derivatives. These phenotypes illustrate that BMP signalling does not control whether cells undergo EMT, but rather ensures that the timing, directionality, and fate outcomes of migrating cells are coordinated with ongoing tissue morphogenesis.

### Implications for EMT-morphogen coordination in evolution and disease

The reciprocal coupling between EMT initiation and BMP pathway competence that we describe establishes a previously unrecognized dimension of EMT function. Beyond its well-characterized roles in breaking down epithelial architecture (e.g. loss of apical-basal polarity, ECM remodelling, cytoskeletal reorganization, and altered cell-cell adhesion), EMT now emerges as an active regulator of morphogen signalling competence. By tuning cellular responsiveness to patterning signals, this additional EMT function provides a mechanism for coordinating migration with fate specification in ways that would be difficult to achieve if these processes were regulated independently.

From an evolutionary perspective, coupling EMT to morphogen responsiveness may represent an important vertebrate innovation for coordinating individual cell behaviours with collective tissue movements. While both EMT machinery and morphogen signalling pathways are ancient and broadly conserved across metazoans (Shook and Keller, 2003; Pérez-Pomares and Muñoz-Chápuli, 2002; Mosby et al., 2024), direct coupling between these processes could have enabled finetuning of invasion timing and coordination, ensuring that precursor region exit and fate specification occurred in concert with tissue-scale movements. The conservation of this coupling across germ layers in the chick embryo suggests it may represent a general developmental principle, though comparative studies across vertebrate species will be needed to assess its evolutionary origins and diversification.

Beyond development, the coupling between EMT and morphogen competence may be relevant for understanding pathological contexts where EMT drives disease progression, particularly cancer metastasis and fibrosis. If similar coupling mechanisms operate in these contexts, therapeutic strategies targeting the reciprocal interactions between EMT state and morphogen responsiveness (rather than either process in isolation) might offer more effective approaches for disrupting pathological invasion. Understanding whether analogous EMT-morphogen couplings exist in other developmental and disease contexts will be important for assessing the generality of this coordination mechanism and its potential as a therapeutic target.

## Resource Availability

### Lead Contact

Requests for further information and resources should be directed to and will be fulfilled by the lead contact, Benjamin Steventon.

## Methods

### EXPERIMENTAL EMBRYOLOGY

#### Fertilized Chicken Eggs

Standard shaver brown fertilized chicken eggs (Medeggs Ltd., Norfolk) and all transgenic chicken eggs (Roslin Institute, Edinburgh) were stored at 18 °C or room temperature (RT) until use and incubated at 37 °C in a humidified incubator (Thermo Fisher Scientific). Depending on the target embryonic stage (Hamburger and Hamilton, 1951), incubation times were specified.

#### Dissecting Embryos

Egg shells were removed using blunt forceps, vitelline membranes just surrounding the embryos were cut using micro-dissecting scissors, and embryos were removed from the egg and transferred into a petri dish with PBS+/+ (Sigma-Aldrich) or saline using a spoon. Under a dissecting microscope (Olympus SZX12), excess yolk on top of the embryos was washed off using glass pasteur pipettes and vitelline membranes were separated from the embryos using sharp forceps. Embryos were visually staged (Hamburger and Hamilton, 1951).

#### Migrating Explants

Migrating explants were prepared as follows (based on methods in Busby et al., 2024). Glass-bottomed dish (MatTek; 35mm Petri dish with 10mm microwell; No. 1.5 coverglass 0.16-0.19mm) microwells were coated with ECM protein of choice (see table below) in PBS+/+ for at least 1h at 37 °C. After removing the ECM protein in PBS+/+ with a pipette, the glassbottom dishes were left to thoroughly dry at 37 °C. All ECM-coated dishes were briefly washed with PBS+/+ twice prior to using. When embryos were finished being dissected, the periphery of the glass-bottom dish wells were covered with silicone grease. 200 *µ*L of warm chick culture media, composed of 10% KnockOut^TM^ Serum Replacement (Thermo Fisher Scientific) + 1% Penicillin-Streptomycin (Thermo Fisher Scientific) + GMEM (Gibco), was added directly to the wells.

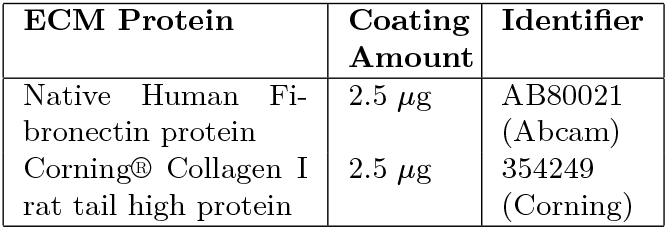

| ECM Protein | Coating Amount | Identifier |
| --- | --- | --- |
| Native Human Fibronectin protein | 2.5 μg | AB80021 (Abcam) |
| Corning® Collagen I rat tail high protein | 2.5 μg | 354249 (Corning) |

Embryonic regions of interest were dissected using a microlance needle (BD) and transferred to the wells using a pipette. Explanted chick tissues were oriented ventral side down and gently flattened using an eyelash knife. Glass coverslips were placed on top to cover the well. Dishes were flooded with PBS+/+ or chick culture media and wrapped with parafilm to prevent dehydration. Explants were either incubated in a tissue culture incubator or live imaged every hour using a widefield microscope (Nikon Eclipse Ti) at 37 °C and 5.5% CO2 for up to 24h.

#### Free Floating Explants

Free floating explants were prepared as follows (based on methods in Streit and Stern, 1999). Embryonic regions of interest were dissected using a microlance needle (BD) and transferred into 24-well plate Millicell® Cell Culture Inserts (Merck) with chick culture media. Plates were incubated in a tissue culture incubator at 37 °C and 5.5% CO2 for up to 24h.

#### Embryo Culture

Embryos were cultured using a modified protocol, based on EC culture methods (Chapman et al., 2001). EC culture plates were prepared by dissolving 0.3g BactoAgar (Sigma-Aldrich) in 50mL of autoclaved 145mM NaCl by heating to boiling, followed by the addition of 50mL pre-warmed thin albumen and penicillin/streptomycin (1:100). The solution was stirred until homogeneous, and 1mL was dispensed into each sterile 35mm Petri dish. Plates were cooled at RT and stored at 4 °C until use. Embryos were transferred to autoclaved filter paper rings and mounted on EC culture. Cultures were placed in a humidified chamber at 37 °C for incubation.

#### Small Molecular Inhibitor Treatment on Embryos and Explants

For whole embryos, inhibitors were added directly to the EC culture agar-albumen. For explants, inhibitors were added directly to the chick culture medium. Inhibitors were used according to the concentration in the table below.

| Inhibitor | Working concentration | Identifier |
| --- | --- | --- |
| LDN193189 hydrochloride | 10nM explants, 100 μM embryos | SML0559 (Sigma-Aldrich) |

#### Electroporation

*Ex ovo* electroporation was performed on chick embryos using an EC culture-based protocol. Embryos were prepared in EC culture and gently cleaned with saline using a mini-pastette (Alpha Laboratories) to remove excess yolk. A plasmid solution was prepared by diluting plasmid DNA (working concentration 1-3 *µ*g/*µ*L) in saline with 0.1% Brilliant Blue dye. Electroporation was conducted using an Intracel TSS20 Ovodyne Electroporator. Embryos were transferred to a custom electroporation chamber filled with sterile saline and positioned over the electrode. The plasmid solution was microinjected into the space between the epiblast and the vitelline membrane using an aspirator tube (Sigma-Aldrich) and pulled 100mm borosilicate capillary tubes with filament (Harvard apparatus). Electroporation parameters for early chick embryos were set to 9.4V, 3 pulses, 50ms duration, and 490ms intervals. Following electroporation, embryos were returned to their EC culture plates and incubated at 37 °C. Plasmid construct expression was typically detectable within 5-6h by fluorescence microscopy or 2-3h by confocal microscopy.

#### Tissue Grafting

Grafting experiments were performed using HH4 stage cytosolic GFP donor embryos and wild-type host embryos. Desired donor tissue were dissected in PBS+/+ and soaked in either recombinant protein or inhibitor solution for 1h at RT (see table below). Grafts were briefly washed in PBS+/+ after inhibitor or protein treatment prior to grafting.

| Inhibitor or Protein | Working concentration | Identifier |
| --- | --- | --- |
| LDN193189 hydrochloride | 1 μM | SML0559 (Sigma-Aldrich) |
| Recombinant Mouse BMP-4 Protein | 10 μg/mL | 5020-BP-010/CF (RD Systems) |

Host embryos were cultured in EC culture and prepared for grafting by cleaning with saline and surgically removing a small region of tissue using an eyelash knife. A plastic ring (cut from an Eppendorf tube) was placed around each host embryo and PBS+/+ was added inside. Treated grafts were transferred using a pipette and carefully placed over the exposed host region using a fluorescent dissecting microscope (Leica). Treated grafts were positioned into the host embryo using an eyelash knife, and excess PBS+/+ was slowly removed while continually monitoring graft position.

#### Tissue Fixation

Whole embryos and tissue explants were fixed in 4% PFA (Sigma-Aldrich) in PBS-/-at RT for 2-3h or at 4 °C overnight.

### MOLECULAR BIOLOGY

#### Bulk RNA Sequencing: Sample Collection

Embryonic and explant tissues (N, biological replicate = 3) from 3-4 dozen embryos each were collected in ice cold PBS+/+.

#### Bulk RNA Sequencing: Extraction & Integrity

Total RNA was extracted using TRIzol^TM^ (Invitrogen) following phase separation and isopropanol precipitation. Samples stored in 1mL TRIzol^TM^ at -80 °C were thawed on ice and homogenized using a cell pestle. Chloroform (200 *µ*L per 1mL TRIzol^TM^) was added, and samples were mixed and incubated at RT for 3min. Following centrifugation at 12,000×g for 15min at 4 °C, the clear aqueous phase was carefully collected, avoiding the interphase, and transferred to a fresh tube. To precipitate RNA, 0.5 *µ*L GlycoBlue (Thermo Fisher Scientific) and 500 *µ*L isopropanol were added to each sample. After 10min at RT, samples were centrifuged at 12,000×g for 10min at 4 °C. The resulting RNA pellet was washed with 1mL of 75% ethanol, vortexed briefly, and centrifuged at 7,500×g for 5min. Ethanol was removed, and pellets were air-dried for 10min before being resuspended in 20 *µ*L RNase-free water and incubated at 55 °C for 10min. RNA integrity was assessed by agarose gel electrophoresis. Approximately 300-500ng of each sample was mixed with loading dye and resolved on a 1% agarose gel in 1× TAE containing GelRed® Nucleic Acid Gel Stain (Biotium). Electrophoresis was carried out at 100V for 1h. Integrity was verified by the presence of distinct 28S and 18S rRNA bands with minimal smearing. Total RNA concentration was measured using the QuBit RNA High Sensitivity kit (ThermoFisher Scientific), and samples were stored at -80 °C until further processing.

#### Bulk RNA Sequencing: Library Prep & Sequencing

Polyadenylated mRNA was enriched from 1 *µ*g of total RNA using the NEBNext Poly(A) mRNA Magnetic Isolation Module (New England Biolabs), following the manufacturer’s guidelines. Libraries were constructed using the NEBNext Ultra II Directional RNA Library Prep Kit for Illumina (New England Biolabs). Dual indexing was performed with NEBNext Multiplex Oligos for Illumina Sets 1 and 2 (New England Biolabs). Cleanup steps throughout library preparation were carried out using SPRIselect magnetic beads (Beckman Coulter). Library quality and fragment size distribution were assessed using an Agilent 2100 Bioanalyzer with High Sensitivity DNA Chips and Reagents (Agilent Technologies). Sequencing was conducted on an Illumina NextSeq 2000 platform (P3 flow cell, singleend 50bp reads, roughly 90 million reads per sample) by Cambridge Genomic Services.

#### Single-cell RNA Sequencing

HH3/3+ embryos were first placed in New culture (New, 1955). Control embryos used in this study were collected as part of a larger node ablation experiment. Embryos were incubated for 0.5h, 2.5h, or 5h following manipulation. Only unablated control embryos were used here; these were detached from the vitelline membrane using an eyelash knife and reattached, to match handling conditions with ablated samples. Embryos were dissected in cold PBS+/+, and approximately half of the area opaca was trimmed to enrich for embryonic tissue. Each embryo was transferred to a 1.5mL Eppendorf tube, washed 3× with PBS+/+, and incubated in 1 mL 0.05% trypsin (Thermo Fisher Scientific) at 37 °C for 15min with gentle trituration every 5min. Trypsin was quenched with 100 *µ*L FBS, and cells were pelleted (450×g, 5min, 4 °C) and resuspended in PBS + 0.05% BSA (Sigma-Aldrich). Cell viability (85–95%) was assessed via haemocytometer. Single-cell suspensions were submitted to the CRUK CI Genomics Core Facility for 10X Genomics library preparation and Illumina sequencing.

#### Immunofluorescence (IF)

Chick embryos and tissue explants were stained following a standard whole-mount immunofluorescence protocol. All washes were performed in PBDT, composed of 1% DMSO (Sigma-Aldrich) + 1% BSA + 1% Triton X-100 (Thermo Fisher Scientific) in PBS-/-, unless stated otherwise. Embryos were blocked for at least 1h at RT in blocking buffer, composed of 2% normal goat serum (Sigma-Aldrich) in PBDT, on a shaker. Primary antibodies were diluted in blocking buffer (see table below) and samples were incubated at 4 °C overnight with gentle agitation. The next day, samples were washed six times for 20min at RT, then incubated overnight at 4 °C in secondary antibody (1:1000) and stains (e.g. DAPI, phalloidin, sytox, all 1:1000) diluted in blocking buffer. Unbound antibody and stains were removed by six 20min washes at RT. Samples were stored in PBDT at 4 °C in the dark until mounting for imaging.

| Antibody | Dilution | Identifier |
| --- | --- | --- |
| Anti slug (C19G7) monoclonal antibody | 1:1000 | #9585 (Cell Signalling Technology) |
| Anti-Smad1 (phospho S463 + S465) antibody | 1:1000 | EPR20662-20 (Abcam) |
| Anti ZO-1 monoclonal antibody | 1:200 | ZO1-1A12 (Thermo Fisher Scientific) |
| Anti PKC zeta monoclonal antibody | 1:200 | sc-17781 (Santa Cruz Biotechnology) |

#### *In Situ* Hybridisation Chain Reaction (HCR)

Fixed samples were washed using PBST, composed of 0.1% Tween 20 (Sigma-Aldrich) in PBS+/+. For longterm storage, samples were dehydrated by a graded series of 20min methanol washes (PBST twice, 25% methanol in PBST, 50% methanol in PBST, 75% methanol in PBST, 100% methanol) and stored at -20 °C. Dehydrated samples were rehydrated in the reverse stepwise manner. *In Situ* detection of mRNA was performed using third-generation hybridization chain reaction (HCR v3.0; Choi et al., 2018). For whole embryos, Proteinase K digestion was carried out by incubating samples in 10 *µ*g/mL Proteinase K (Thermo Fisher Scientific) in PBST for either 3min (stages HH8 and younger) or 5min (stages HH8 and older) at RT. Samples were then post-fixed in 4% PFA in PBS-/-for 20min at RT. Proteinase K digestion and post-fixation were omitted for explant samples. All samples were washed with PBST for 5min twice, followed by a 5min wash in 5×SSCT (Thermo Fisher Scientific; diluted in ultrapure water) at RT. Samples were then incubated at 37 °C for 30min in probe hybridization buffer (Molecular Instruments) for pre-hybridization. To prepare the probe solution, 2 *µ*L of the 1 *µ*M stock of each probe set was added per 500 *µ*L of hybridization buffer, and samples were incubated at 37 °C overnight. On the following day, samples were washed four times in pre-warmed HCR wash buffer (Molecular Instruments) at 37 °C for 15min, then washed twice in 5×SSCT at RT for 5min. To equilibrate, samples were incubated in amplification buffer (Molecular Instruments) at RT for at least 5min. Hairpins were snap-cooled individually by heating 30pmol of each (from a 3 *µ*M stock) to 95 °C for 90 seconds in a thermocycler and allowing them to cool to RT in the dark. Equal volumes of snap-cooled hairpins H1 and H2 were combined in amplification buffer, and samples were incubated at RT overnight, protected from light. On the last day, samples were washed thrice in 5×SSCT for 15min at RT. Nuclear staining was performed by incubating samples in 5×SSCT containing DAPI (1:1000) for 30min at RT, followed by three 10min washes in 5×SSCT. Samples were stored in 5×SSCT at 4 °C in the dark until mounting for imaging.

#### Mounting

Stained samples were mounted between two coverslips with VECTASHIELD® Antifade Mounting Medium (Vector Laboratories).

#### Gelatin Embedding Stained Embryos

IF- and HCR-stained embryos were embedded in gelatin for cryosectioning. Stained embryos were protected from light at all steps. First, stained embryos were washed with PBS-/-. Stained embryos were then cryoprotected by incubation in 5% sucrose (Thermo Fisher Scientific) in PBS-/-at RT for 10-30min, followed by overnight incubation in 15% sucrose in PBS-/-at 4 °C until fully sunk. The next day, embryos were equilibrated in molten gelatin (7.5% gelatin + 15% sucrose + PBS-/-) at 37 °C for 3-5h, then embedded in cryomolds (Agar Scientific). Embryos were positioned near the bottom of the mold using forceps, and blocks were solidified at 4 °C for at least 5min. Once set, molds were flash-frozen by brief sequential submersion in liquid nitrogen and stored at -80 °C until sectioning.

#### Cryosectioning

Gelatin-embedded embryos were cryosectioned using a Microtome Leica Cryostat 3050S. Samples were equilibrated inside the cryostat chamber (chamber temperature -20 °C; object temperature -14 °C) for at least 10min prior to sectioning. Sample blocks were mounted onto the specimen stage using Epredia^TM^ Neg-50^TM^ Frozen Section Medium (Thermo Fisher Scientific) or OCT compound (Agar Scientific), and trimmed at 50 *µ*m before sectioning at 25 *µ*m. Sections were collected onto Epredia^TM^ SuperFrost Plus^TM^ Adhesion slides (Thermo Fisher Scientific) and air-dried overnight at RT in the dark. Slides were stored at 4 °C, protected from light, until imaging.

#### *SNAI2* Overexpression

The coding sequence of *SNAI2*, spanning from the ATG start codon to the stop codon, was amplified from cDNA by PCR with flanking XhoI restriction sites. The amplified fragment was first cloned into the pSC-A plasmid (Strataclone) for sequence verification. Subsequently, the verified insert was subcloned into the pCIG expression vector using XhoI, positioned upstream of the IRES2.

#### *SNAI2* CRISPR-Cas9

CRISPR-Cas9-mediated gene editing was performed as previously described (Williams et al., 2018). The Cas9-expressing plasmid pCAG Cas9-2A-Citrine (Addgene #92358) and the chick-specific sgRNA vector pcU6-3 sgRNA (Addgene #92359) were used for electroporation. *SNAI2* sgRNA target sequences were selected using CHOPCHOP (Labun et al., 2019) based on low off-target potential and the presence of a diagnostic restriction enzyme site for genotyping. sgRNAs were cloned into the pcU6-3 vector using annealed oligonucleotides and Golden Gate assembly with BsmBI, followed by transformation, colony PCR, and sequencing to confirm correct insertion. Genomic DNA was extracted from dissected tissues using either crude extraction of alkaline lysis (50mM NaOH, 95 °C) followed by neutralization (1M Tris-HCl, pH 8.0) or PureLink^TM^ Genomic DNA Mini Kit (Invitrogen^TM^). Genotyping was performed by PCR amplification of the target region, restriction digest, and agarose gel electrophoresis to distinguish wild-type from successfully CRISPR-edited mosaic tissues.

### DATA ANALYSIS

#### Bulk RNA Sequencing Data Analysis

Raw reads were processed using the Galaxy web platform (Afgan et al., 2016). Quality control was performed using FastQC (Simon Andrews et al., 2010), adaptor trimming was performed with Trim Galore! (Krueger et al., 2023), alignment to the galGal5 reference genome from the Genome Reference Consortium was carried out using HISAT 2 (Kim et al., 2015), PCR duplicates were removed using SAMtools markdup (Li et al., 2009), transcript assembly was performed with StringTie (Pertea et al., 2015), and differential expression analysis was carried out using DESeq2 (Love et al., 2014).

#### Gene Ontology

Gene ontology enrichment analyses were conducted using GeneCodis 4 (Garcia-Moreno et al., 2022).

#### Single-cell RNA Sequencing Analysis

CellRanger (10X Genomics) filtered feature matrices were passed into Seurat (Satija et al., 2015) in R v4.2.3. Reads were mapped to the galGal6 reference genome from the Genome Reference Consortium. The following quality control steps were taken: checking the % of mitochondrial genes, filtering out cells with too low or high number of different genes expressed, normalising the data, and regressing for both sex effect and cell cycle. Principal Component Analysis (PCA) was then performed for dimensionality reduction, followed by Uniform Manifold Approximation and Projection (UMAP) using the first 15 principal components (dims = 1:15) and a clustering resolution of 0.2 for visualization of cellular transcriptomic profiles. Feature plots were generated to display the expression patterns of genes of interest across the UMAP space. Correlation analyses were conducted between the expression levels of genes of interest across individual cells to assess coexpression dynamics.

#### IMAGING & IMAGE ANALYSIS

Fixed and HCR- or immuno-stained samples were imaged on the Zeiss LSM700 confocal microscope. Images were analysed using FIJI, to create either maximum projection or resliced images. Snapshots of live embryos on EC culture plates were imaged on a fluorescent dissecting scope (Leica) in brightfield and GFP channels. Images were overlaid using FIJI. Live imaging of embryos on EC culture plates and explants were performed using Nikon Eclipse Ti widefield microscope. Further quantification of HCR- or immuno-stained samples were performed using IMARIS.

## Contributions

Y.T. and B.S. conceived the project, designed experiments, and wrote the manuscript. Y.T. performed all experiments and analysed the data. A.N. and L.B. performed the scRNA-seq experiment. F.T. contributed to electroporation and LDN culture experiments. C.C. designed and made the *SNAI2* overexpression construct. G.S.N. contributed to code used in analyses.

## Acknowledgments

We thank Felipe Karam Teixeira, Fran Dearden, Nozomi Takahashi, Tim Grocott, Kathy Niakan, and past and present members of the Xiong, Scarpa, Santos, Clark, Rosello-Diez, and Steventon labs for technical assistance and constructive feedback on the project. We thank Kevin Costello for assistance in analysing the bulk RNA sequencing data. We thank Jon Howe and Ben Sutcliffe from the University of Cambridge Microscopy Bioscience Platform Light Microscopy Team for microscopy support. Funding: this work was supported by a Herchel Smith PhD Studentship and Research Support grant to Y.T. and a Wellcome Trust Discovery Award (225360_Z_22_Z) to B.J.S.

## Declaration of Interests

The authors declare no competing interests.

## Declaration of generative AI and AI-assisted technologies in the writing process

During the preparation of this work, the author(s) used Claude in order to proofread (checking for spelling errors, reducing word count, and checking for typos) this manuscript. After using this tool/service, the author(s) reviewed and edited the content as needed and take(s) full responsibility for the content of the published article.

## Supplementary Figures

**Fig. S1.**
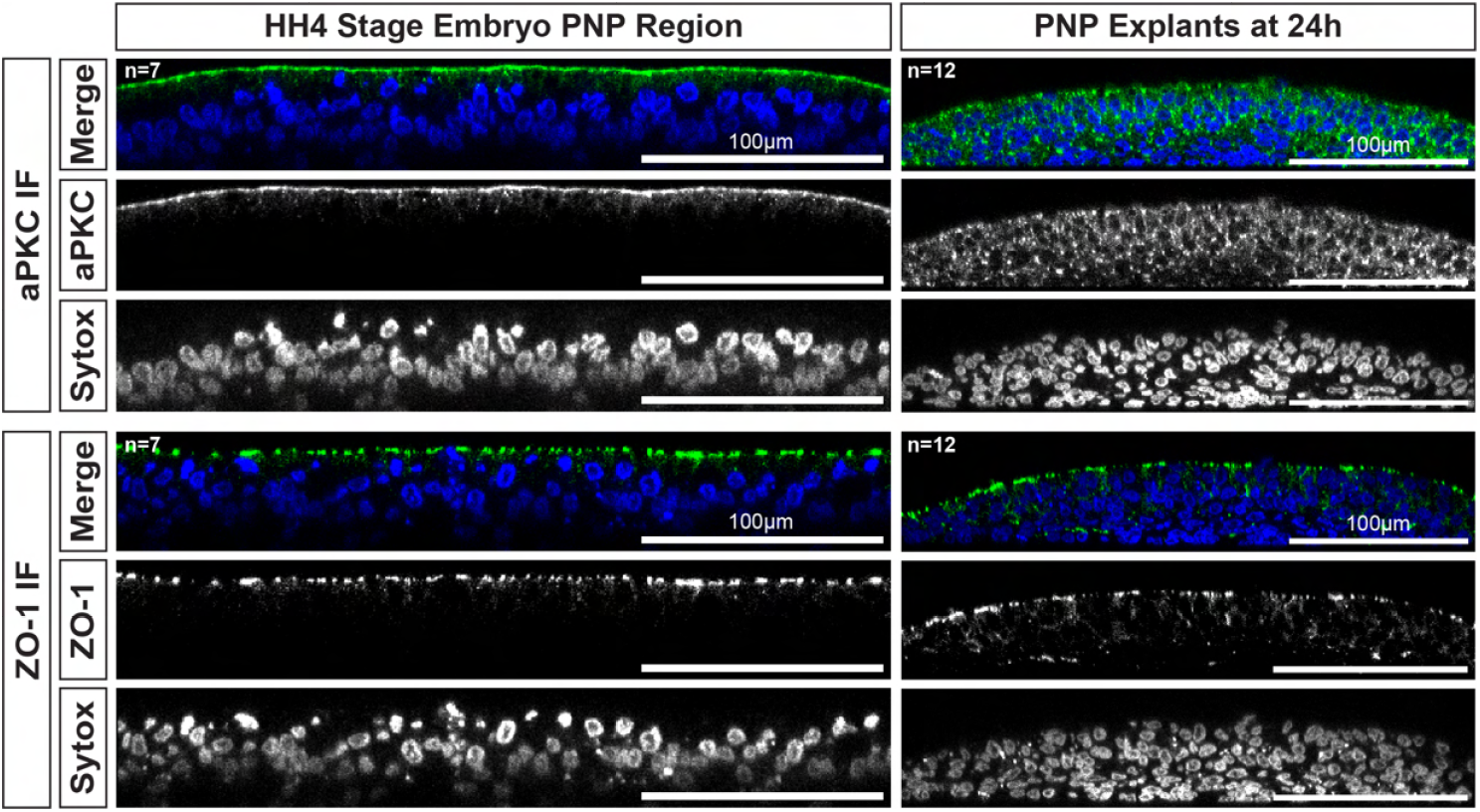
Loss of apical basal polarity observed in PNP migrating explants. Reslices of representative confocal image of aPKC and ZO-1 immunofluorescence and SYTOX staining on the PNP embryonic region and PNP migrating explants on human fibronectin after 24h of incubation.

**Fig. S2.**
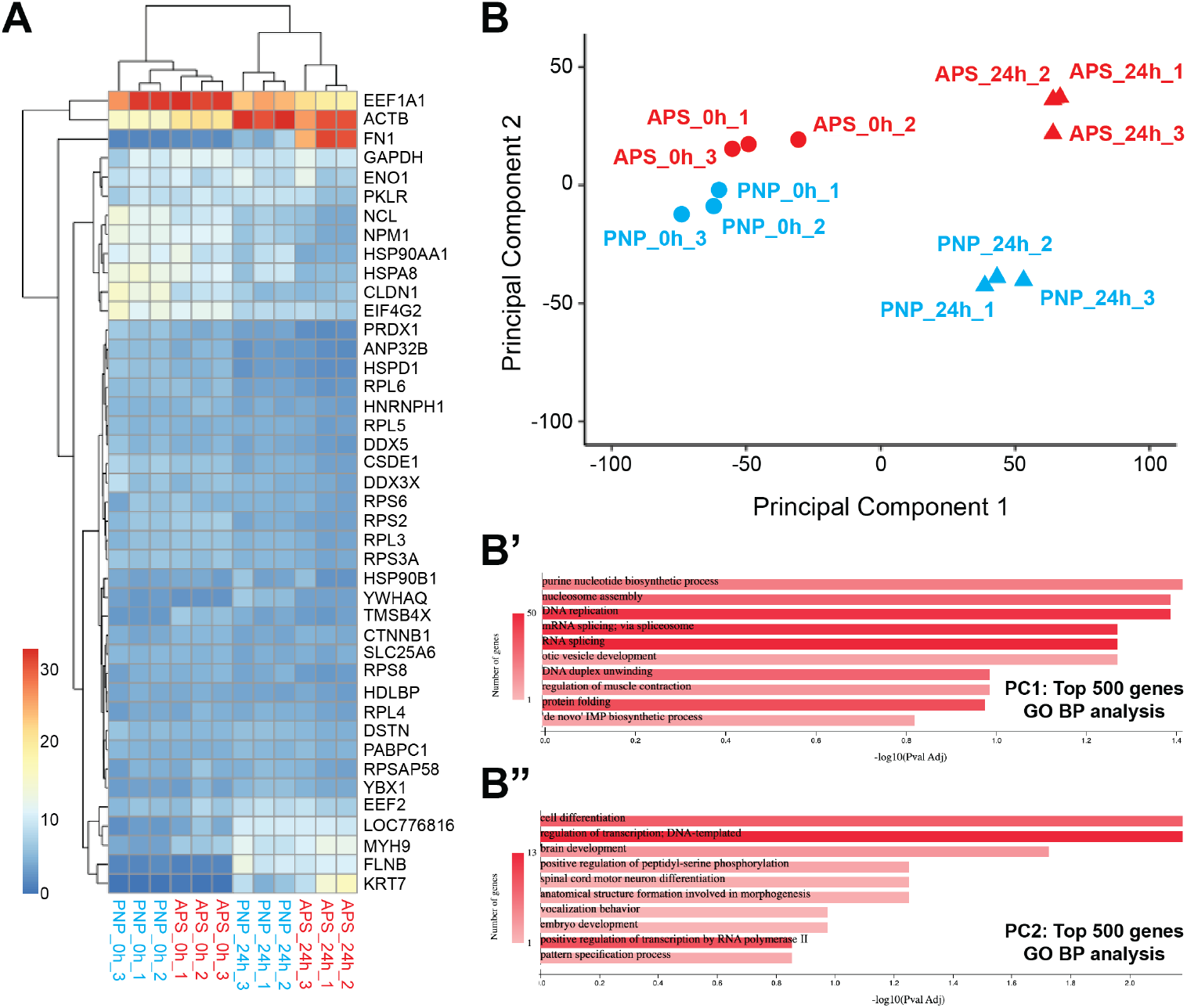
Bulk RNA sequencing reveals common genes upregulated or downregulated in migrating explants, regardless of tissue type. A) Z-score heatmap showing the top differentially expressed genes across all samples, filtered by a summed z-score threshold (>50). Genes are hierarchically clustered, and samples are grouped by tissue type (PNP vs APS) and timepoint (24h vs 0h). B) Principal component analysis (PCA) of all bulk RNA sequencing samples reveals clear separation of samples at 24h vs 0h along PC1 and PC2. PC1 accounts for variation primarily associated with DNA replication, transcription, and translation. PC2 corresponds to changes in cell differentiation status. B’) Gene Ontology (GO) Biological Process (BP) enrichment of the top 500 genes contributing most to PC1, highlighting pathways associated with biosynthesis, translation, and cell cycle progression. B”) GO BP enrichment of the top 500 genes contributing most to PC2, showing enrichment for terms related to differentiation and developmental processes.

**Fig. S3.**
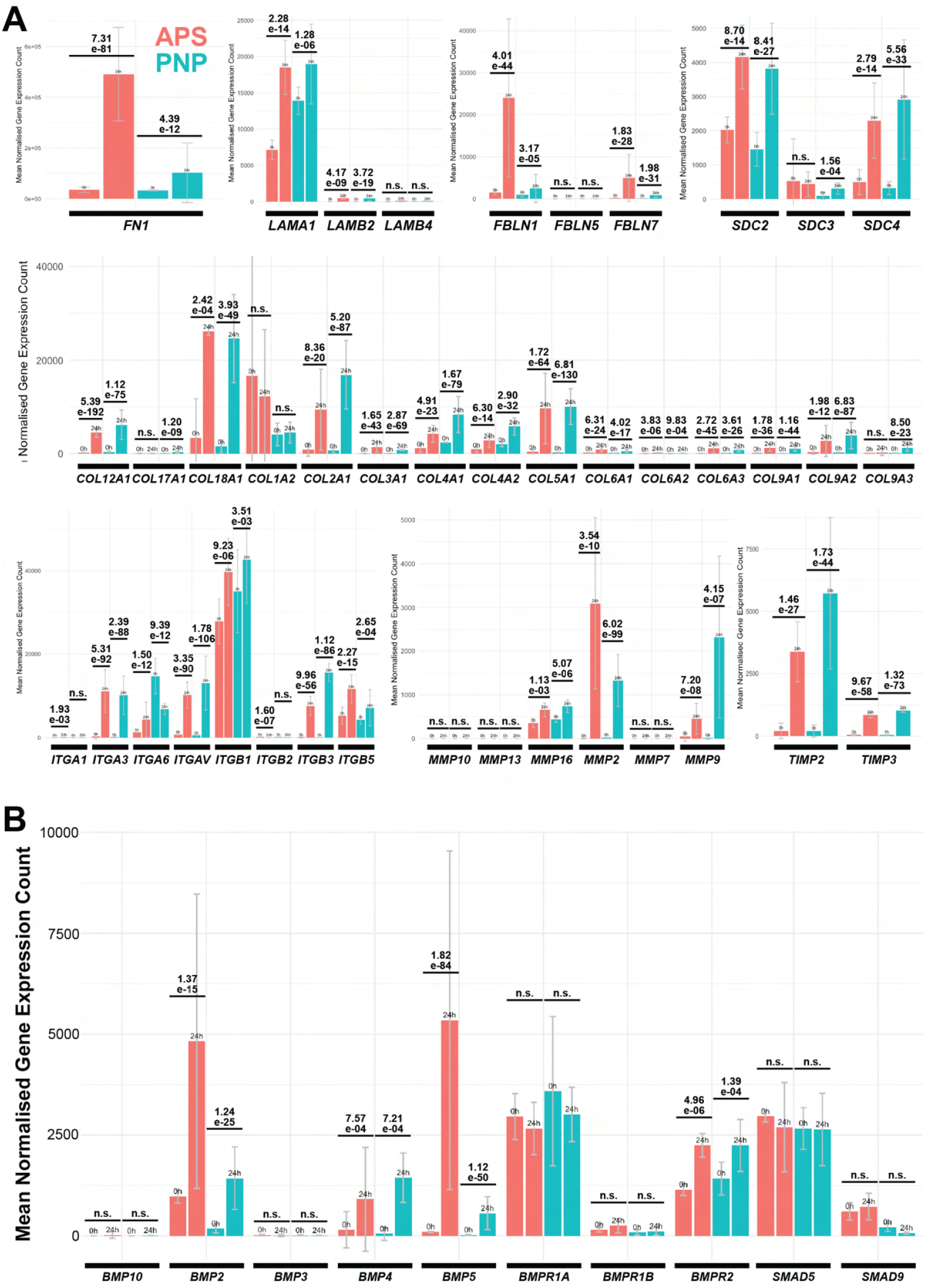
Bulk RNA sequencing reveals upregulation of many ECM components, ECM remodelling enzymes, and BMP signalling pathway components. A) Bulk RNA sequencing mean normalised gene expression counts for ECM component and ECM remodelling genes of interest in PNP and APS migrating explants at 0h vs 24h of incubation. FN: Fibronectin. LAM: Laminin. FBLN: Fibulin. SDC: Syndecan. COL: Collagen. ITG: Integrin. MMP: Matrix metalloproteases. TIMP: Tissue inhibitor of metalloproteinases. B) Bulk RNA sequencing mean normalised gene expression counts for genes encoding BMP ligands and receptors in PNP and APS migrating explants at 0h vs 24h of incubation.

**Fig. S4.**
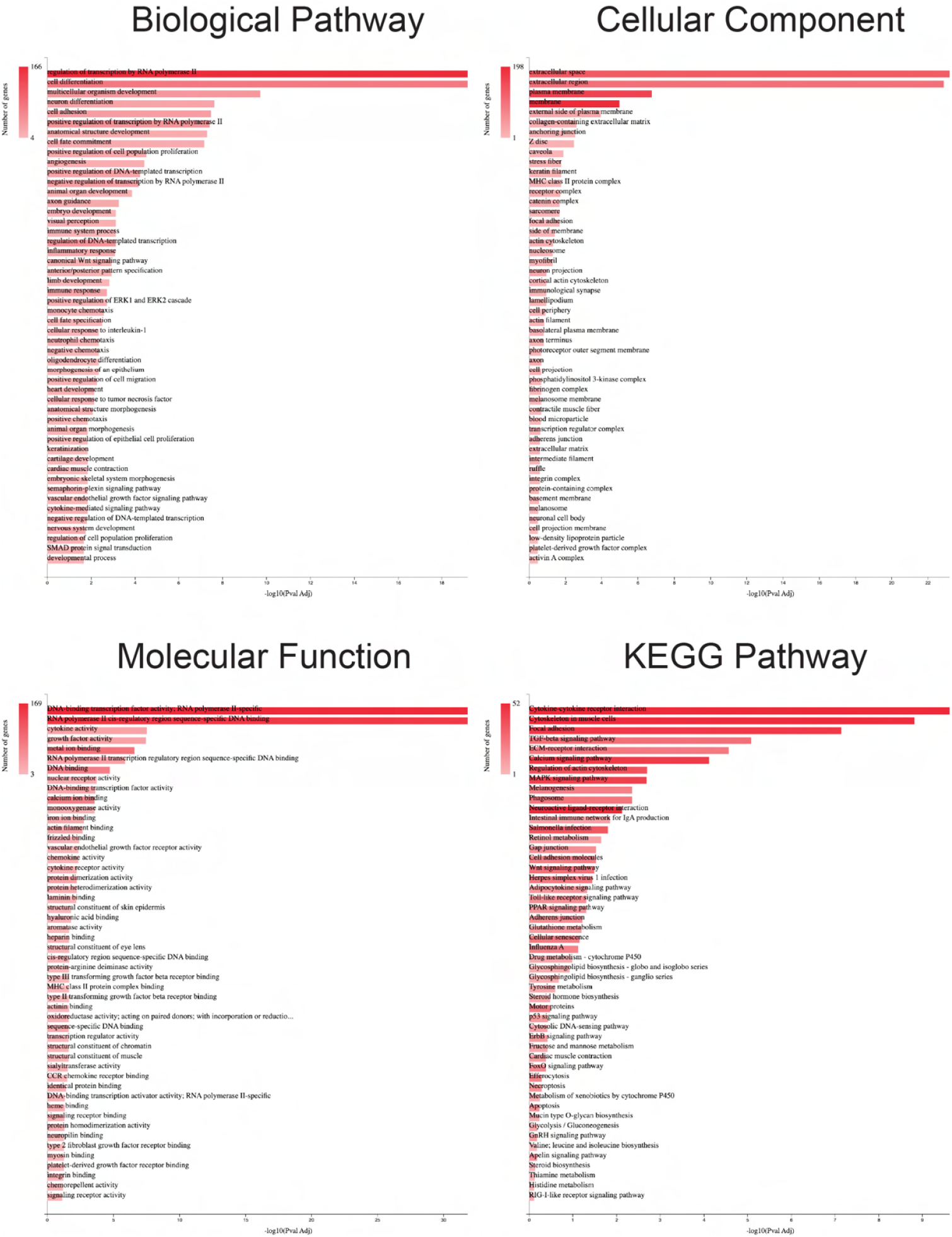
Gene ontology of genes upregulated at least 2-fold in both PNP and APS tissue upon migration. Top 50 terms shown.

**Fig. S5.**
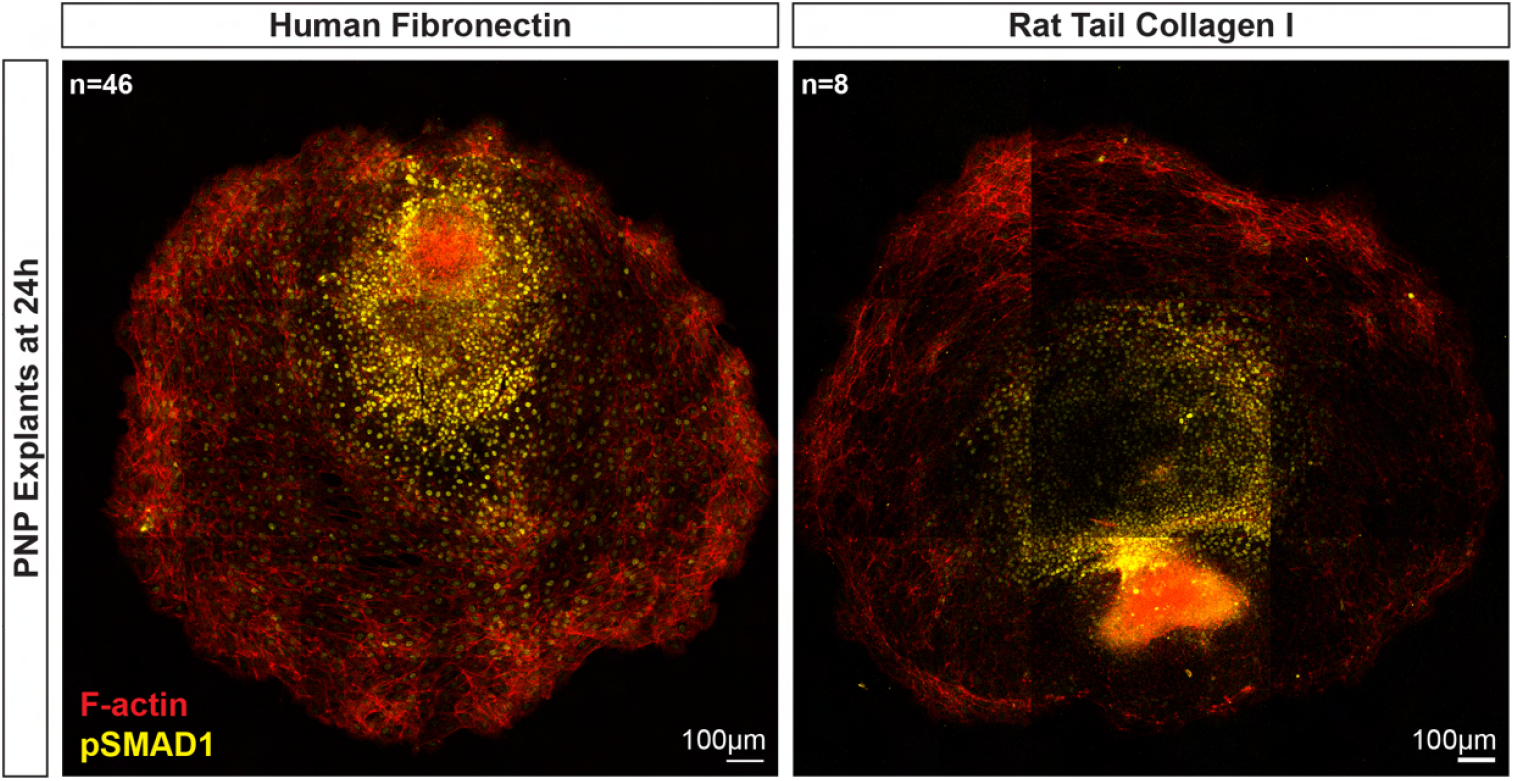
BMP signalling activity induced in PNP *ex vivo* migrating explants on fibronectin and collagen. Representative confocal image of phospho-SMAD1 immunofluorescence and F-actin staining on an PNP migrating explant on human fibronectin vs. rat tail collagen I after 24h of incubation.

**Fig. S6.**
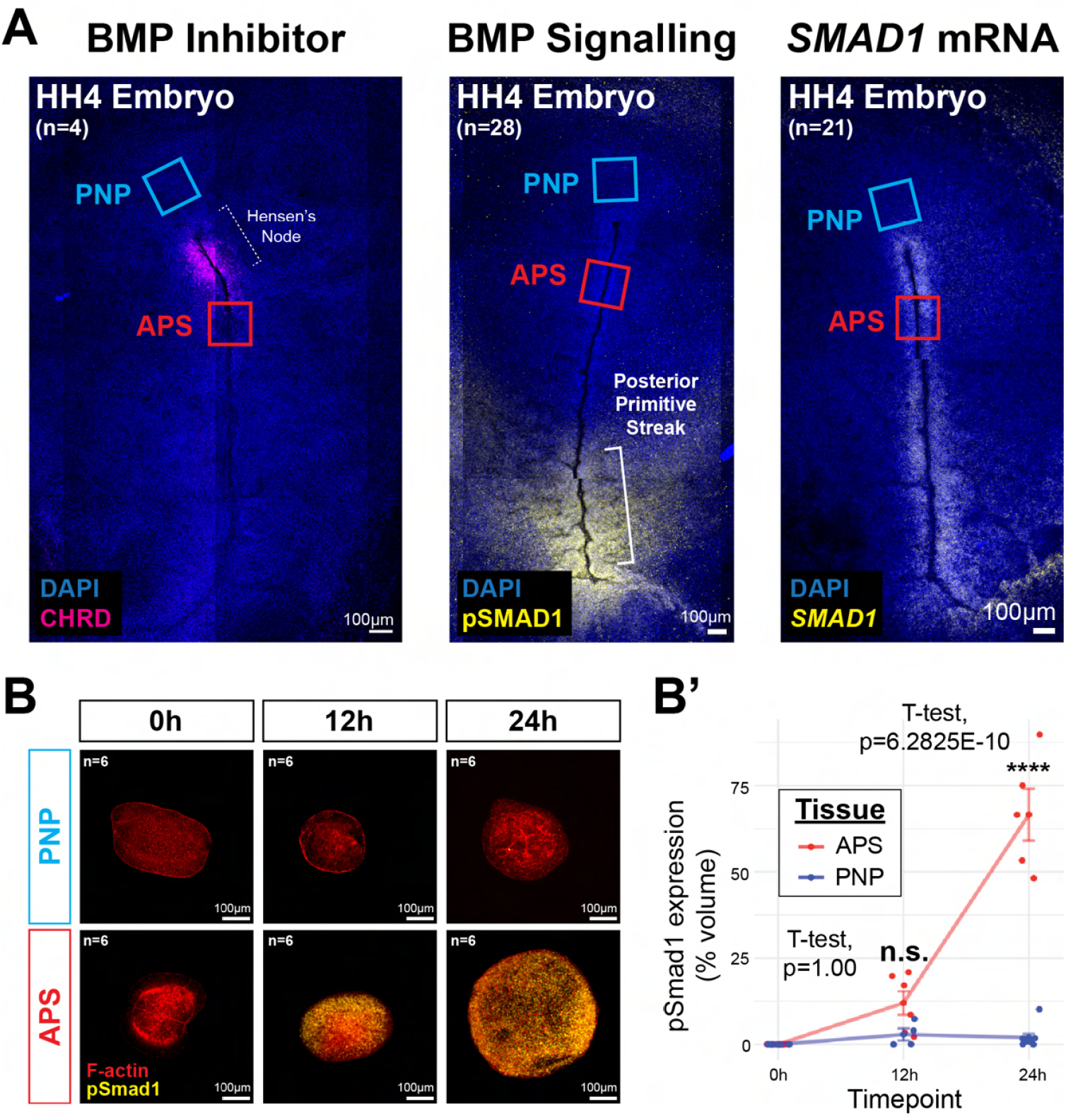
BMP signalling activation in PNP migrating explants is not due to lifting of BMP inhibition by Hensen’s node. A) Representative confocal images of HH4 stage chick embryo *CHRD in situ* HCR staining (left), phospho-SMAD1 immunostaining (middle), and *SMAD1 in situ* HCR staining (right). B) Representative pseudo-time course confocal images of phospho-SMAD1 immunofluorescence and F-actin stain in PNP and APS floating explants. B’) Quantification of phospho-SMAD1 expression as percent volume of sytox signal in PNP and APS floating explants.

**Fig. S7.**
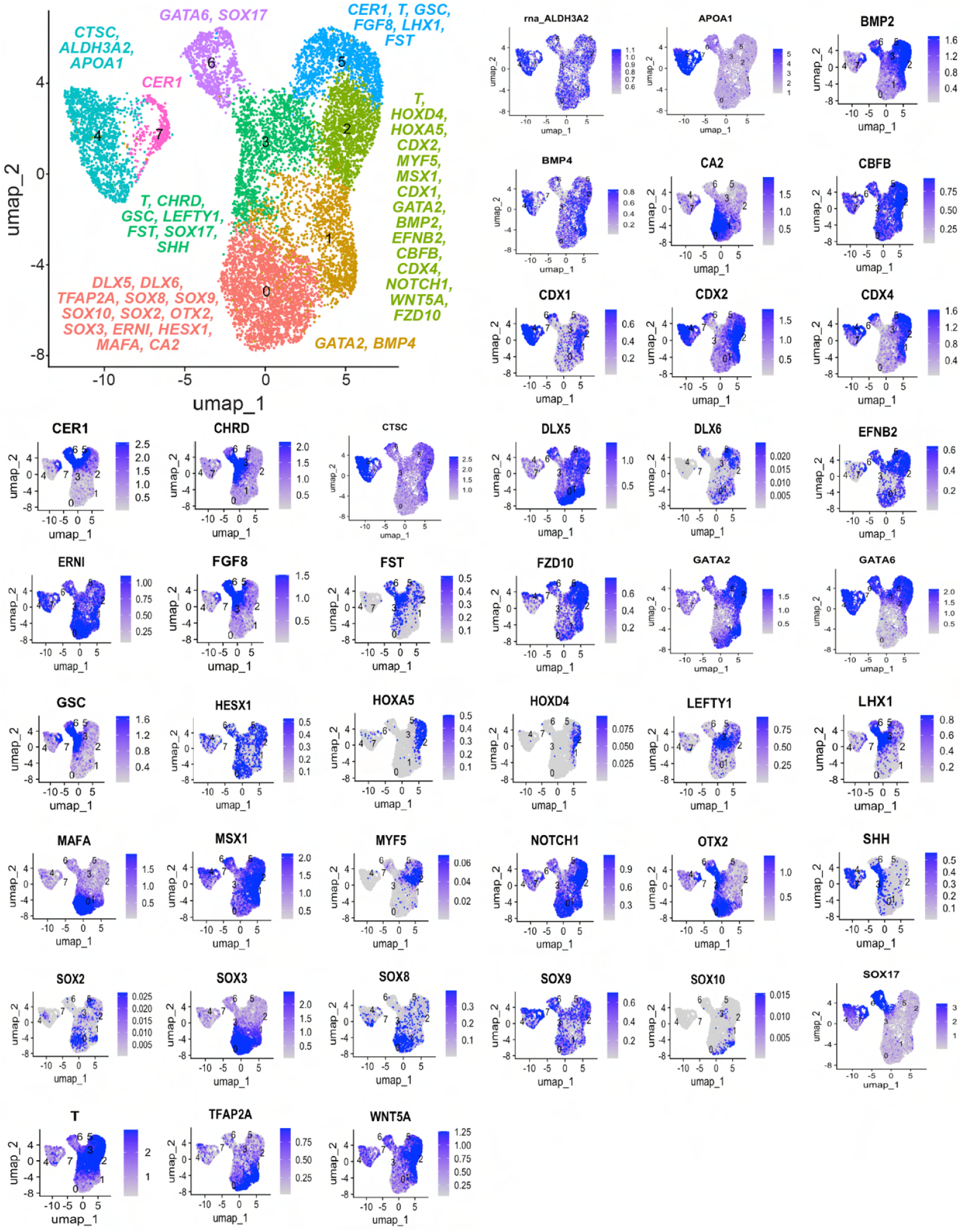
UMAP cluster annotations. A List of relevant top differentially expressed genes per cluster, and their feature maps in alphabetical order.

**Fig. S8.**
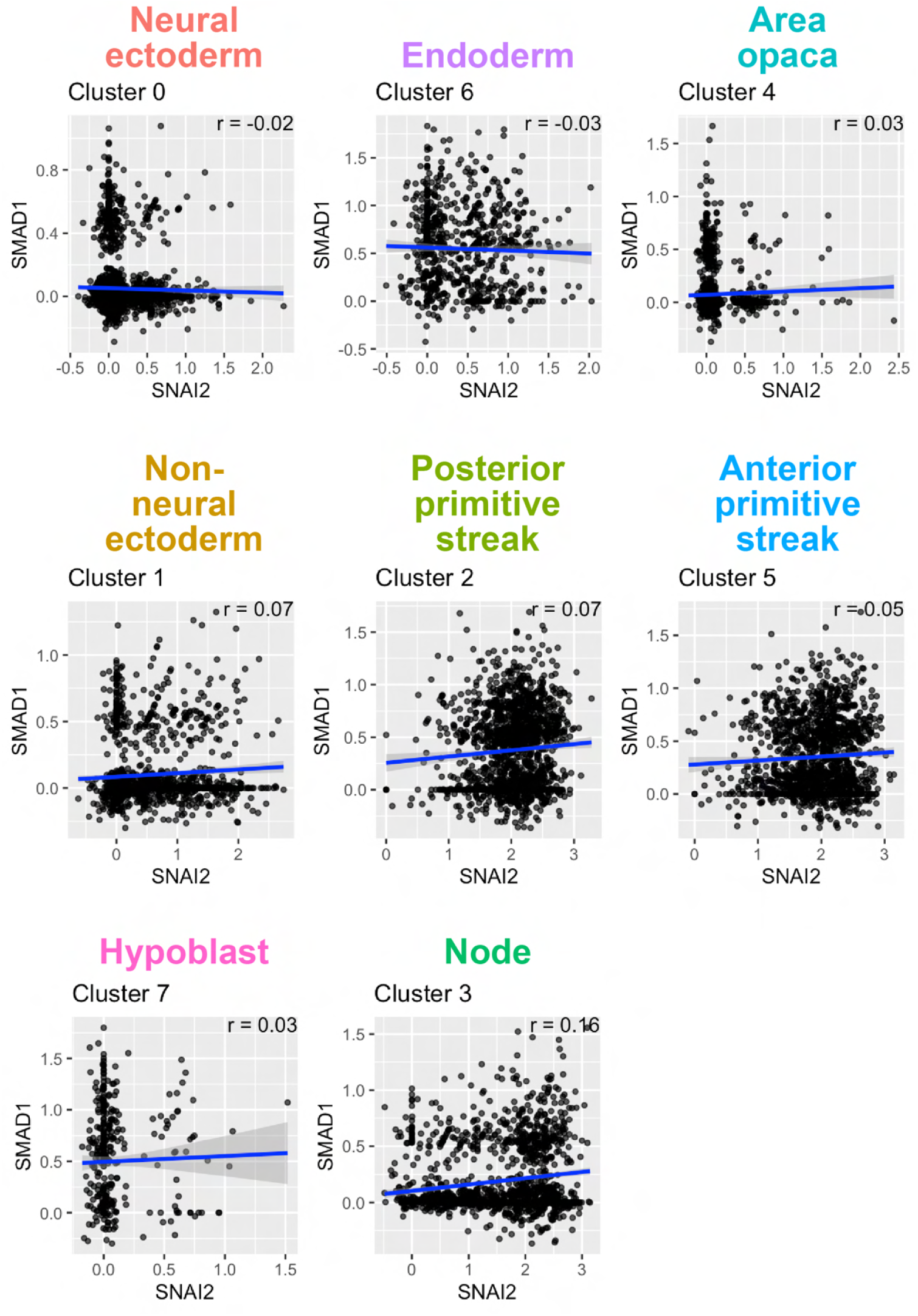
Correlation between *SNAI2* and *SMAD1* gene expression in individual clusters. Pearson’s correlation between *SMAD1* and *SNAI2* expression.

**Fig. S9.**
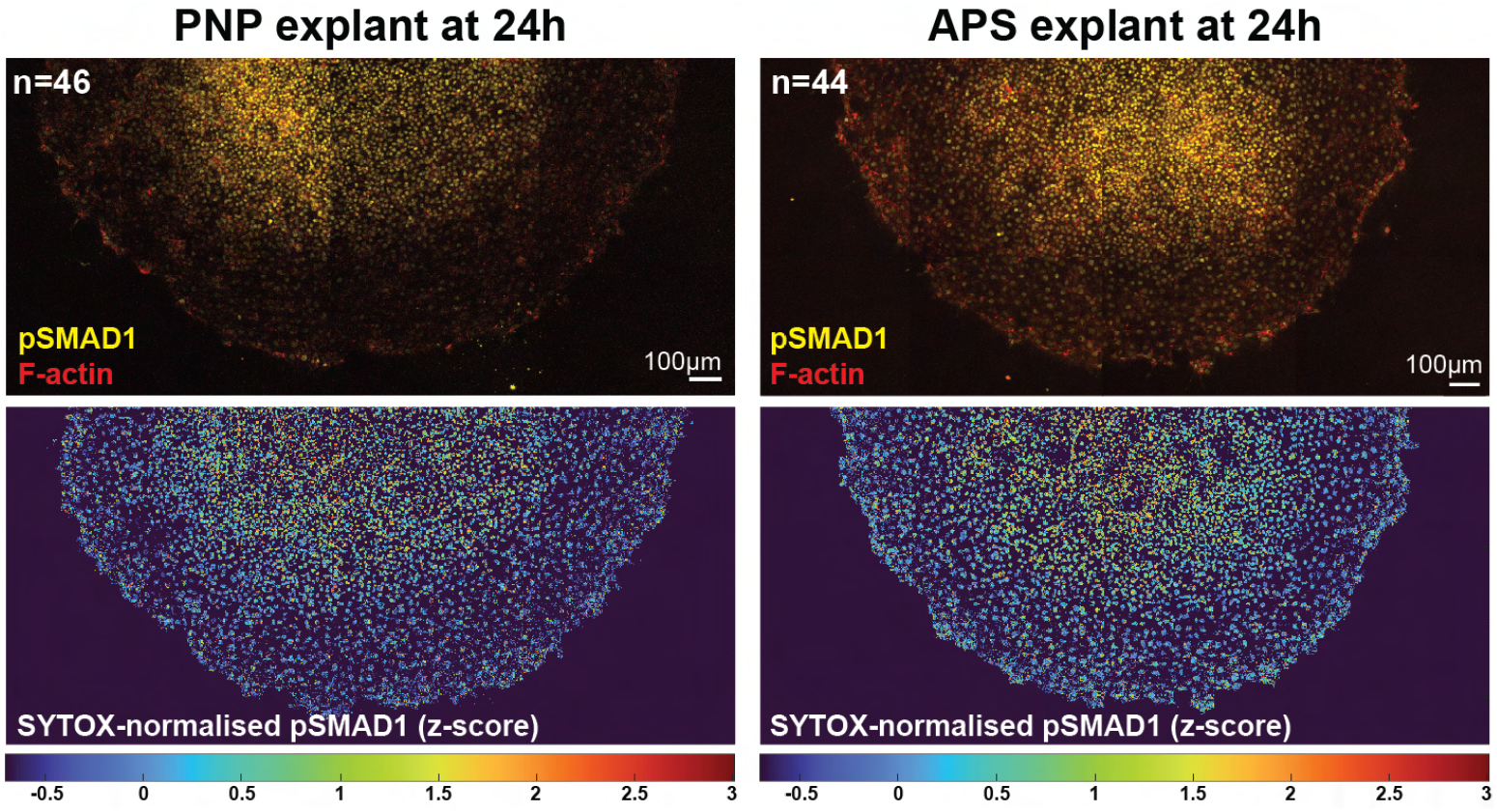
BMP signalling patterns in *ex vivo* explants. Top: Representative confocal images of pSMAD1 immunostaining and F-actin staining in PNP and APS explants on human fibronectin after 24h of incubation. Bottom: Z-scores of SYTOX-normalised pSMAD1 signal intensities. Z-scores tend to decrease towards the migrating edges.

**Fig. S10.**
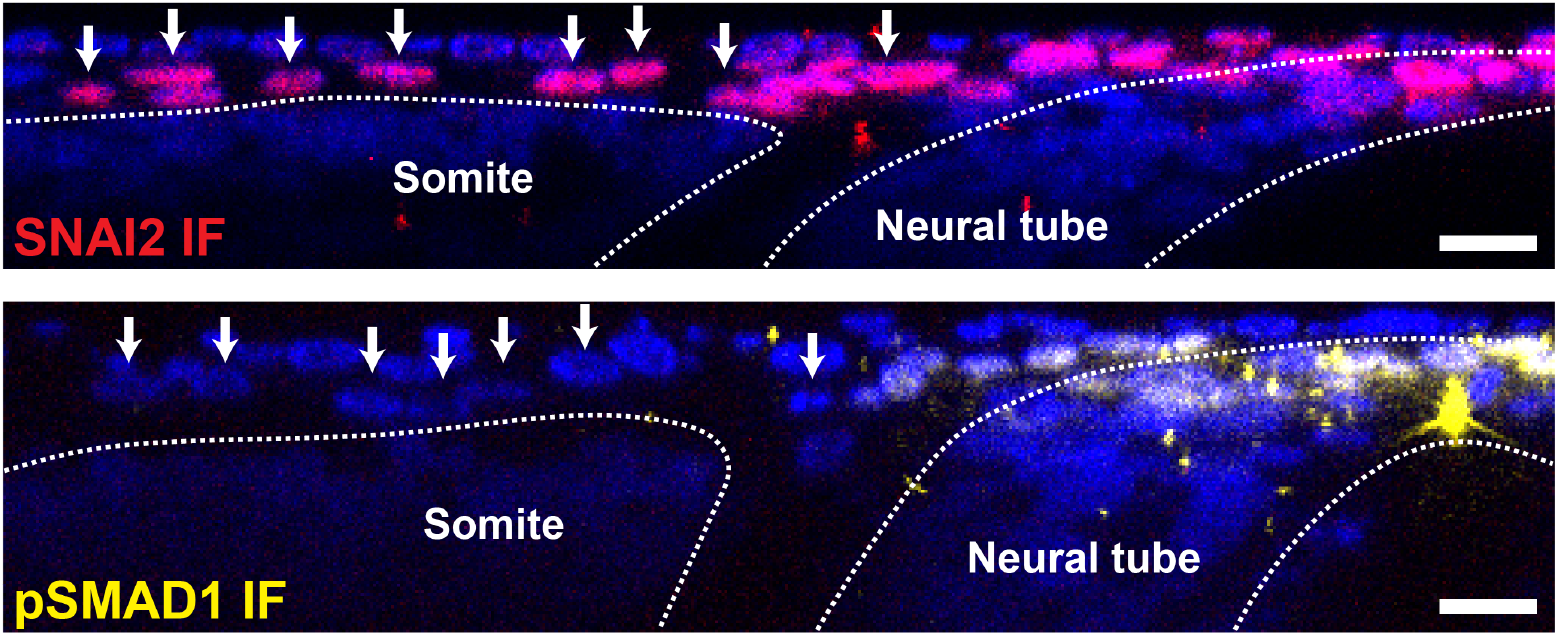
Reslices of dorsal neural tube and migrating neural crest cells. Top: Reslice of representative SNAI2 immunostaining. Bottom: Reslice of representative pSMAD1 immunostaining. White dotted lines show somite and neural tube boundaries.

**Fig. S11.**
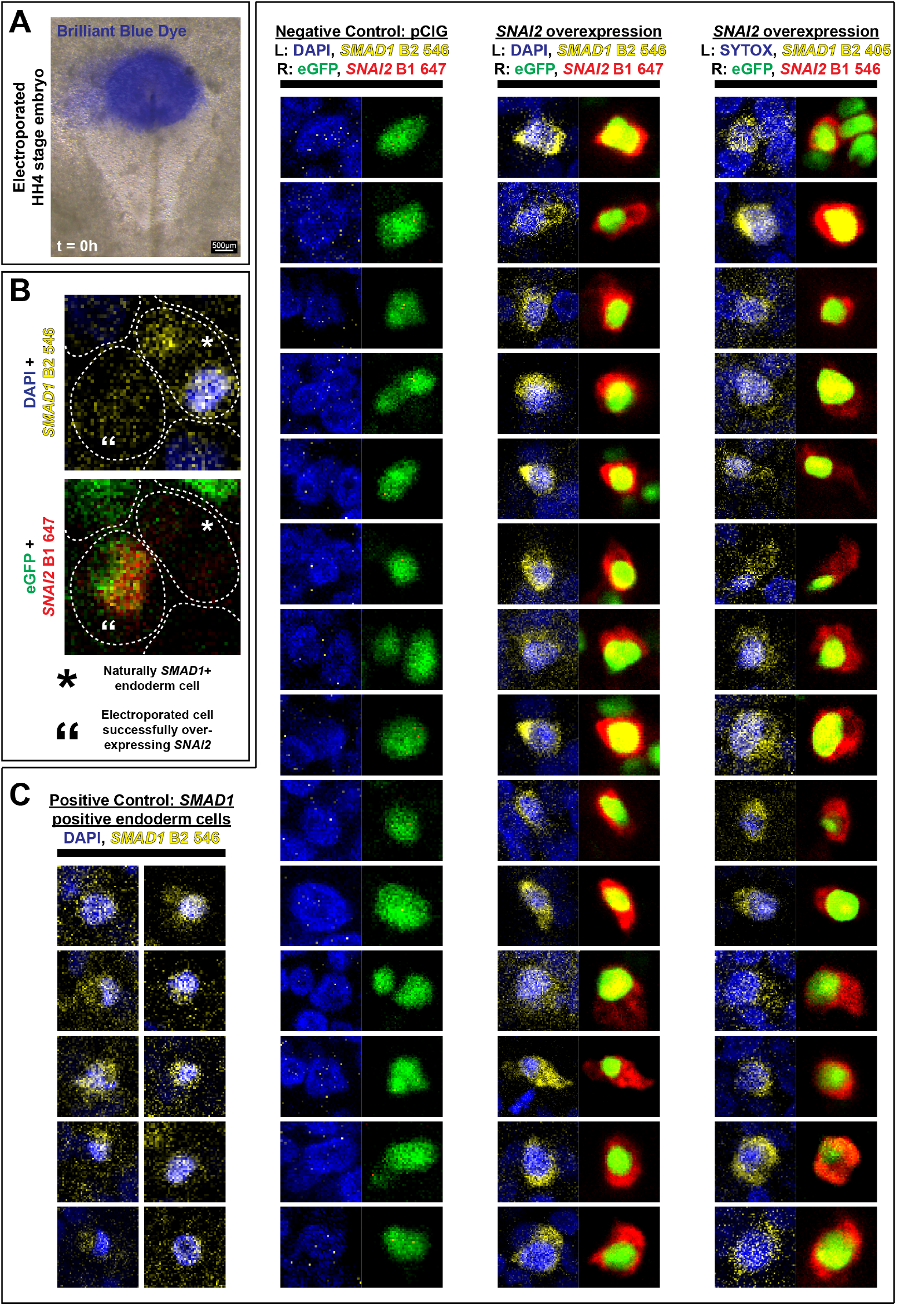
*SNAI2* overexpression is sufficient to drive *SMAD1* expression. A) Representative brightfield image of an electroporated embryo at t=0h. B) A representative confocal image showing a naturally *SMAD1* -positive endoderm cell next to an electroporated cell. C) Representative confocal images of *SNAI2* and *SMAD1 in situ* HCR staining of endoderm cells (left; positive control for *SMAD1* expression), pCIG-electroporated cells (middle; negative control for *SNAI2* expression), and cells electroporated with the *SNAI2* -overexpression construct (right two columns; experimental; different HCR hairpin combinations).

**Fig. S12.**
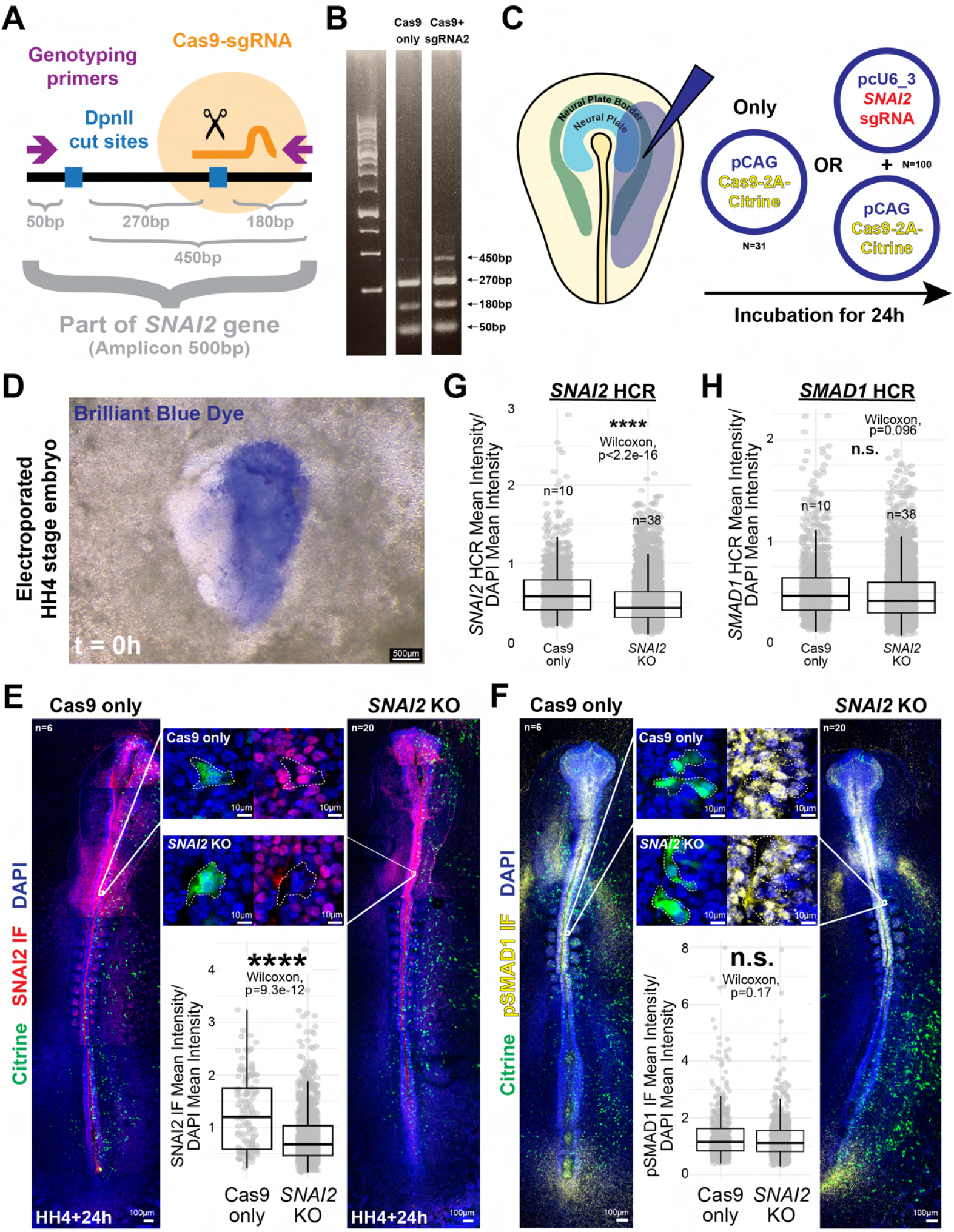
Insufficient knock out of *SNAI2* to test necessity hypothesis. A) Schematic of sgRNA design. B) Gel image showing an extra 450bp band in only the mosaic knockout lane. C) Schematic of CRISPR-Cas9 *SNAI2* knockout experiment in HH4 neural plate border. Embryos electroporated with pCAG Cas9-2A-Citrine (Cas9 expression vector) alone or with pcU6-3 *SNAI2* -sgRNA (*SNAI2* sgRNA expressing vector) and incubated for 24h. D) Representative brightfield image of an electroporated embryo at t=0h. E) SNAI2 immunofluorescence in pCAG Cas9-2A-Citrine vs pCAG Cas9-2A-Citrine + pcU6-3 *SNAI2* -sgRNA electroporated embryos. IMARIS quantification of SNAI2 immunofluorescence signal intensities normalised by DAPI signal intensities in Citrine-positive cells. F) pSMAD1 immunofluorescence in pCAG Cas9-2A-Citrine vs pCAG Cas9-2A-Citrine + pcU6-3 *SNAI2* -sgRNA electroporated embryos. IMARIS quantification of pSMAD1 immunofluorescence signal intensities normalised by DAPI signal intensities in Citrine-positive cells. G) *SNAI2 in situ* HCR signal quantification in Citrine-positive cells. H) *SMAD1 in situ* HCR signal quantification in Citrine-positive cells.

**Fig. S13.**
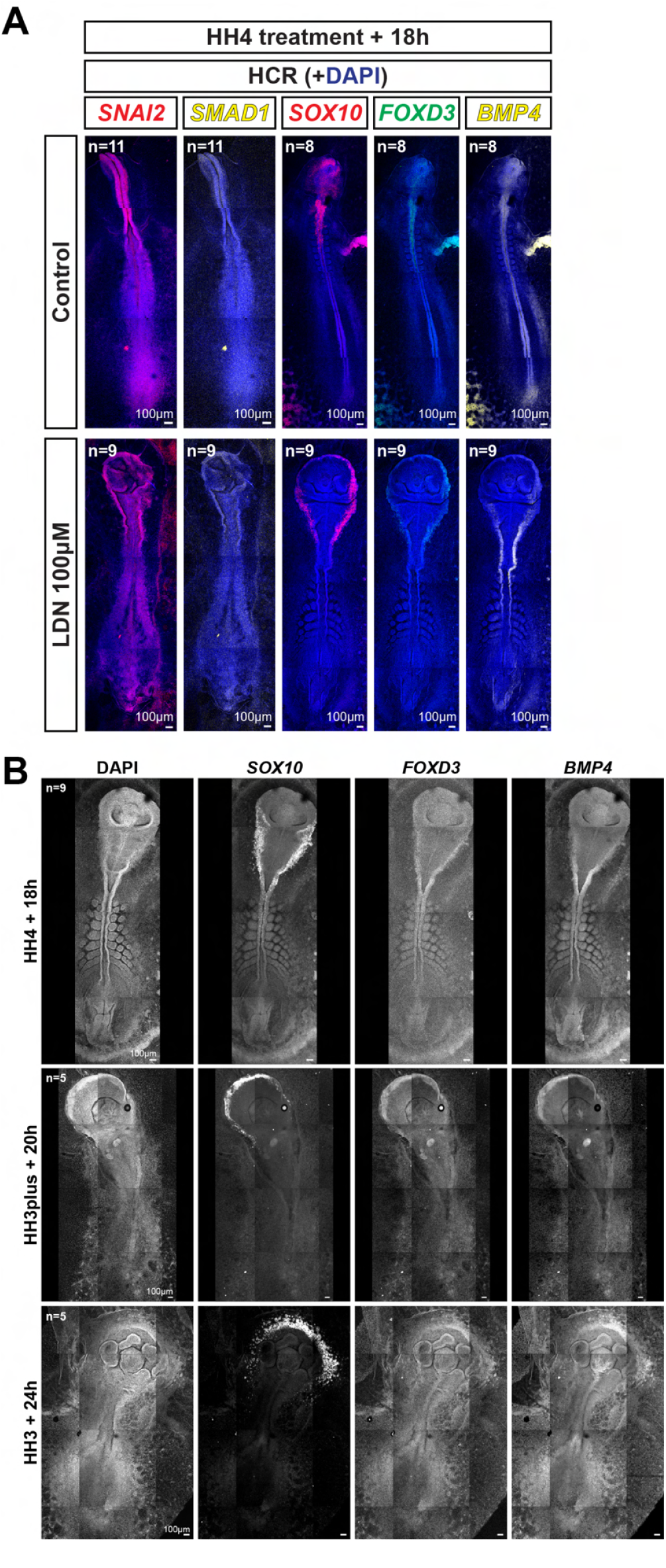
Neural crest markers are still expressed in LDN-treated embryos. A) Representative confocal images of *SNAI2* /*SMAD1* and *SOX10* /*FOXD3* /*BMP4 in situ* HCR staining in control vs LDN-treated embryos. B) Representative greyscale confocal images of *SOX10* /*FOXD3* /*BMP4 in situ* HCR and DAPI staining of chick embryos treated with LDN from HH4 stage (top), HH3+ stage (middle), or HH3 stage (bottom).

**Fig. S14.**
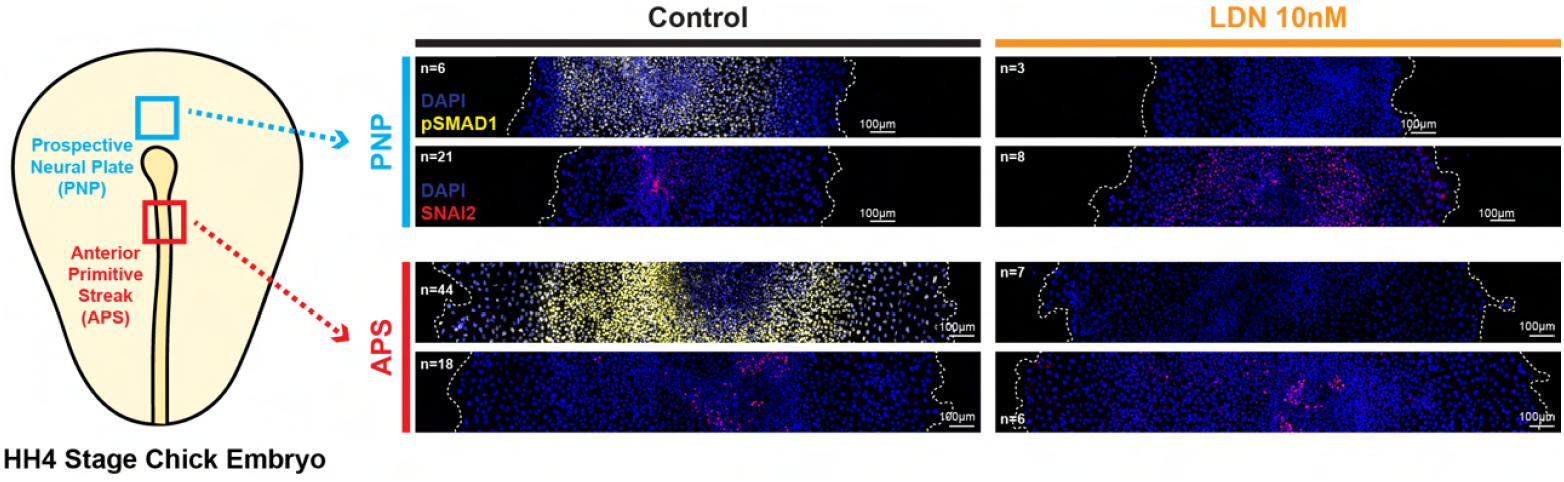
BMP signalling not needed for EMT initiation in *ex vivo* explants. Representative confocal images of pSMAD1 and SNAI2 immunofluorescence and SYTOX staining in control and LDN-treated PNP and APS explants on human fibronectin after 24h of incubation.

**Fig. S15.**
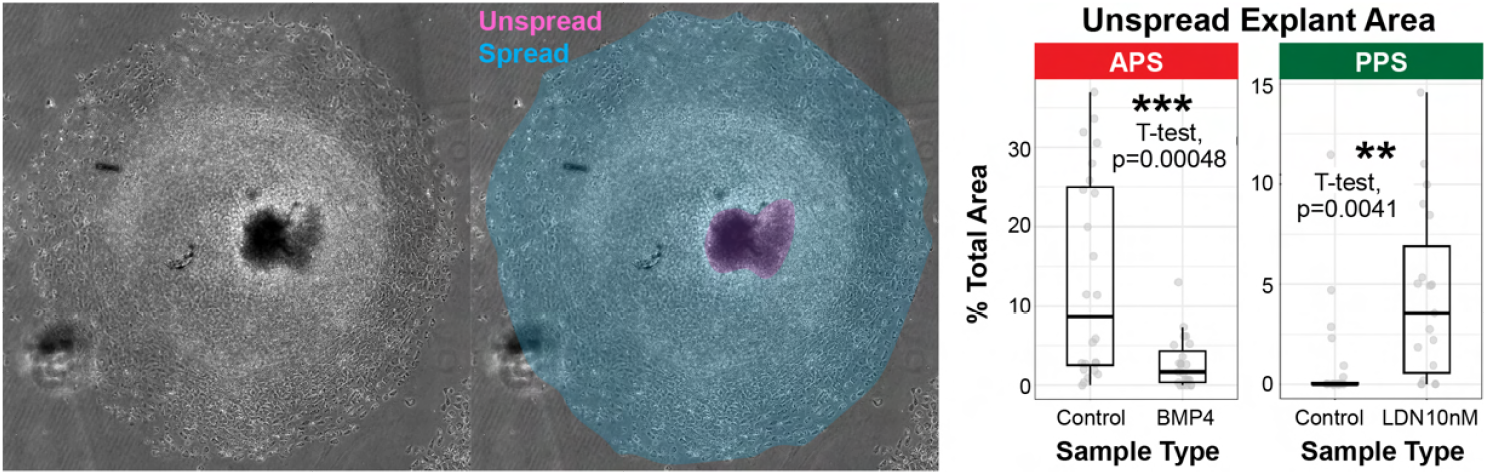
BMP signalling controls the extent of spreading in *ex vivo* explants. Left: Representative spread vs. unspread areas. Right: Quantification of explant area still left unspread at final time point.

**Fig. S16.**
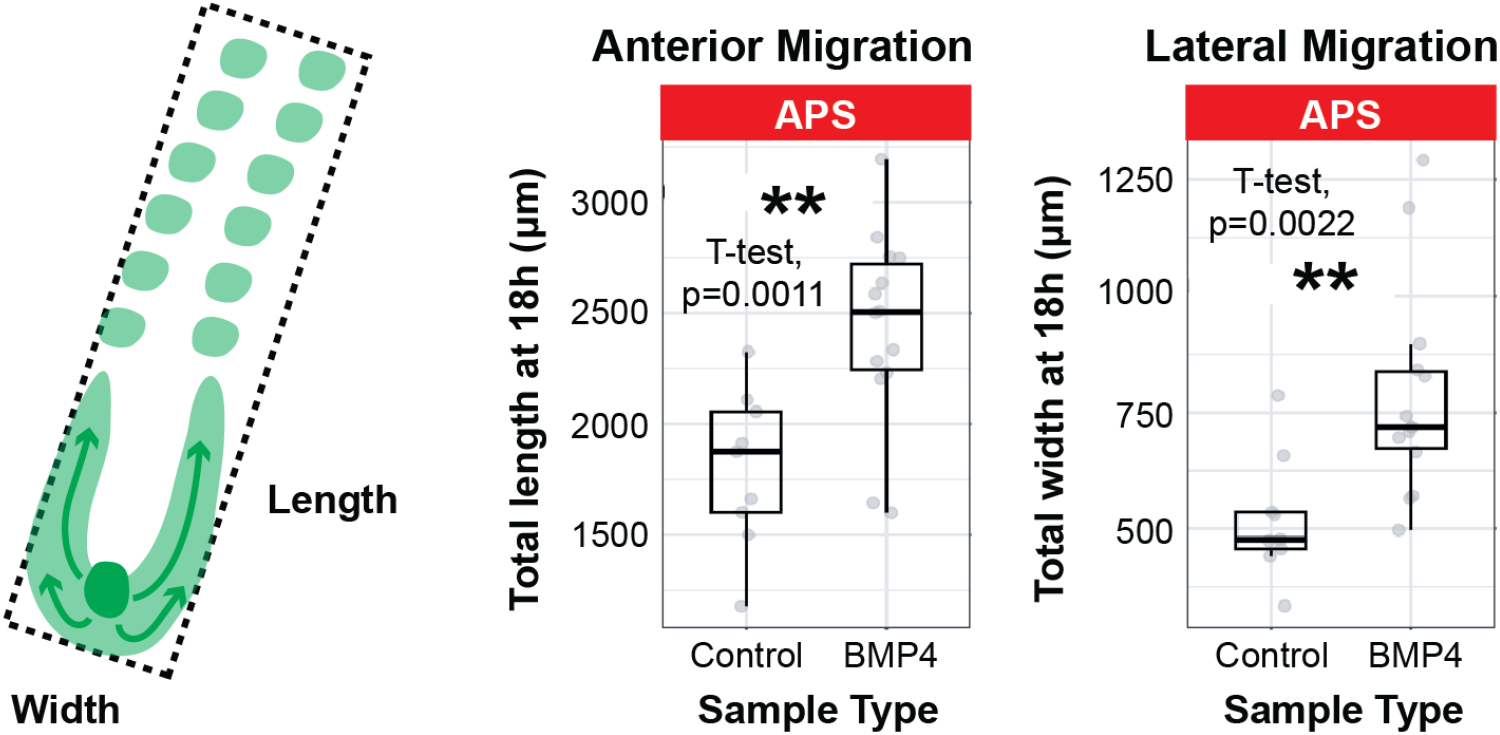
BMP4-treated APS grafts contribute more anteriorly and laterally compared to controls. Left: Schematic of graft length and width. Right: Quantification of APS graft anterior and lateral migration.

## Notes

### Competing Interest Statement

The authors have declared no competing interest.

